# Steady-state neuron-predominant LINE-1 encoded ORF1p protein and LINE-1 RNA increase with aging in the mouse and human brain

**DOI:** 10.1101/2023.12.12.571308

**Authors:** Tom Bonnifet, Sandra Sinnassamy, Olivia Massiani-Beaudoin, Philippe Mailly, Héloïse Monnet, Damarys Loew, Berangère Lombard, Nicolas Servant, Rajiv L. Joshi, Julia Fuchs

**Affiliations:** CIRB, Collège de France, Université PSL, INSERM, CNRS, 75005, Paris, France; Orion Technological Core, CIRB, Collège de France, Université PSL, INSERM, CNRS, 75005, Paris, France; Institut Curie, Université PSL, Centre de Recherche, CurieCoreTech Spectrométrie de Masse Protéomique, 26 rue d’Ulm, Paris 75248 Cedex 05, France; Institut Curie, INSERM U900, Mines Paris Tech, Université PSL

**Keywords:** Transposable elements, retrotransposons, LINE-1, ORF1p, aging, brain mapping, mass spectrometry, ORF1p protein interactors, human post-mortem brain, dopaminergic neurons, deep learning

## Abstract

Recent studies have established a reciprocal causal link between aging and the activation of transposable elements, characterized in particular by a de-repression of LINE-1 retrotransposons. These LINE-1 elements represent 21% of the human genome, but only a minority of these sequences retain the coding potential essential for their mobility. LINE-1 encoded proteins can induce cell toxicity implicated in aging and neurodegenerative diseases. However, our knowledge of the expression and localization of LINE-1-encoded proteins in the central nervous system is limited. Using a novel approach combining atlas-based brain mapping with deep-learning algorithms on large-scale pyramidal brain images, we unveil a heterogeneous, neuron-predominant and widespread ORF1p expression throughout the murine brain at steady-state. In aged mice, ORF1p expression increases significantly which is corroborated in human post-mortem dopaminergic neurons by an increase in young LINE-1 elements including those with open reading frames. Mass spectrometry analysis of endogenous mouse ORF1p revealed novel, neuron-specific protein interactors. These findings contribute to a comprehensive description of the dynamics of LINE-1 and ORF1p expression in the brain at steady-state and in aging and provide insights on ORF1p protein interactions in the brain.

## Introduction

Only about 2 % of the human genome are DNA sequences that will be translated into protein. The remaining 98 % are comprised of introns, regulatory elements, non-coding RNA, pseudogenes and repetitive elements including transposable elements. However, some sequences in what is generally considered “non-coding genome” do in fact contain sequences which encode proteins. This is true for specific lncRNAs which can encode peptides or functional proteins ^1^ but also for a few copies of two transposable element families, Long INterspersed Element-1 (LINE-1) and Human Endogenous RetroViruses (HERV). Non-functional copies of retrotransposons, to which LINE-1 and HERV belong, cover about 44% ^2^ of the human genome as remnants of an evolutionary ancient activity. Depending on the source, about 100 ^3^ to 146 ^4^ full-length LINE-1 elements with two open reading frames encoding ORF1 and ORF2 are present in the Human reference genome (GRCh38 Genome Assembly) and several incomplete HERV sequences encoding either or any combination of envelope (env), gag, pro or pol ^5^. The LINE-1 encoded protein ORF1p, an RNA binding protein with “cis” preference ^6, 7^, and ORF2p, an endonuclease and reverse transcriptase ^8, 9^ are required for the mobility of LINE-1 elements. As many other transposable elements (TEs) including HERVs, LINE-1 elements are repressed by multiple cellular pathways. It was thus generally thought that TEs are repressed in somatic cells with no expression at steady-state ^10–12^. However, the aging process reduces the reliability of these repressive mechanisms ^13^. It is now, 31 years after the initial proposition of the “transposon theory of aging” by Driver and McKechnie ^14^, still a matter of debate whether TE activation can be both, a cause and a consequence of aging ^15, 16^.

Sparse data has shown that the LINE-1 encoded protein ORF1p is expressed at steady-state in the mouse ventral midbrain ^17^, the mouse hippocampus ^18^ and in some regions of the human post-mortem brain ^19^ and recent data informed about the presence of full-length transcripts in cancer cells, human epithelial cells and mouse hippocampal neurons ^20^. Repression of LINE-1 might thus be incomplete and if so, it remains unclear how cells then prevent cell toxicity associated with LINE-1 encoded protein activity. Indeed, LINE-1 encoded proteins have been demonstrated to induce genomic instability (ORF2p endonuclease-mediated ^17, 21–26)^ and inflammation (ORF2p reverse transcriptase-mediated ^27–29)^ and these cellular activities might be causally related to organismal aging, cancer, autoimmune and neurological diseases ^30^. For instance, LINE-1 activation can drive neurodegeneration of mouse dopaminergic neurons ^17^, of drosophila neurons ^31, 32^ and of mouse Purkinje neurons ^33^ which can be at least partially rescued with nucleoside analogue reverse transcriptase inhibitors (NRTIs) or other anti-LINE-1 strategies. NRTIs are currently being tested in several clinical trials designed to target either the RT of HERVs or the RT encoded by the LINE-1 ORF2 protein. It is not known today, however, to which extend LINE-1 encoded proteins are expressed at steady-state throughout the mouse and human brain, whether there is cell-type specificity and whether activation of LINE-1 encoded proteins is associated with brain aging or human neurodegeneration. Here, using a deep-learning assisted cellular detection methodology applied to pyramidal large-scale images of the mouse brain mapped to the Allen mouse brain atlas combined with post-mortem human brain imaging, co-IP mass spectrometry and transcriptomic analysis of LINE-1 expression, we describe a brain-wide map of ORF1p expression and interacting proteins at steady-state and in the context of aging. We find a heterogenous but widespread expression of ORF1p in the mouse brain with predominant expression in neurons. In aged mice, neuronal ORF1p expression increases brain-wide and in some brain regions to up to 27%. In human dopaminergic neurons, young LINE-1 transcripts and specific full-length and coding LINE-1 copies are increased in aged individuals. We further describe endogenous mouse ORF1p interacting proteins revealing known interactors and unexpected interacting proteins belonging to GO categories related to RNA metabolism, chromatin remodeling, cytoskeleton and the synapse.

## Results

### Widespread and heterogenous expression of the LINE-1 encoded ORF1p protein in the wildtype mouse brain

To investigate the expression pattern and intensities of endogenous LINE-1 encoded ORF1p protein throughout the entire mouse brain, we devised a deep-learning assisted cellular detection methodology applied to pyramidal large-scale images using a comprehensive workflow complemented by an approach based on confocal imaging as schematized in Figure 1A. Briefly, starting from sagittal slide scanner images of the mouse brain, we defined anatomical brain regions by mapping the Allen Brain Atlas onto the slide scanner images using Aligning Big Brains & Atlases (ABBA) ^34^. We then employed a deep-learning detection method to identify all cell nuclei (Hoechst) and categorize all detected cells into neuronal cells (NeuN+) or non-neuronal cells (NeuN-) and ORF1p-expressing cells (ORF1p+) or cells that do not express ORF1p (ORF1p-). This workflow allowed us then to characterize the cell identity of ORF1p+ cells and ORF1p intensity throughout the whole brain but also in specific anatomical regions. In parallel, we completed the approach using confocal microscopy on selected anatomical regions allowing for comparison with higher resolution. Importantly, the specificity of the ORF1p antibody, a widely used, commercially available antibody ^18, 35–39^, was confirmed by blocking the ORF1p antibody with purified mouse ORF1p protein resulting in the complete absence of immunofluorescence staining (Suppl Fig. 1A), by using an in-house antibody against mouse ORF1p^17^ which colocalized with the anti-ORF1p antibody used (Suppl Fig. 1B, quantified in Suppl Fig. 1C), by immunoprecipitation and mass spectrometry used in this study (Fig 6A, Suppl Table 2) and by siRNA-mediated knock-down of ORF1 in a differentiated mouse dopaminergic cell line (MN9D; Suppl Fig. 1D). Unexpectedly, we found a generalized and widespread expression of ORF1p throughout the brain of wildtype mice (Fig. 1B; Swiss OF1 mice, three months-old; whole brain except regions with particularly high cellular density (cerebellum, hippocampus, olfactory bulb) which impedes nuclei detection by deep-learning. ORF1p is detectable in all regions and subregions analyzed with heterogenous expression patterns (density and intensity) per region/subregions. The ten regions shown in Figure 1B exemplify visible different densities of ORF1p+ cells with varying levels of expression. Notably, the expression pattern of ORF1p in the hippocampus is similar to what has recently been published ^18^ (Fig. 1B, panel 2). Throughout the entire brain, the mean density of ORF1p+ cells per mm² was ≈ 305 ±18 (mean ±SEM), representing up to 20% of all detected cells (Fig. 1C). ORF1p+ cells in each mouse brain analyzed showed up to eight-fold disparities in intensity between low- and high-expressed cells (Fig. 1C). We then quantified nine anatomical regions according to the Allen Brain Atlas on four brains of three-month old mice (Fig. 1D) using the automated workflow (Fig. 1A) with regard to cell density (Fig. 1D), cell proportions (Fig. 1E) and fluorescent intensity of ORF1p+ cells (Fig. 1F). This approach permitted the analysis of about 10 000 ORF1p+ cells per animal highlighting the power of our large-scale analysis. Densities of ORF1p+ cells ranged from the lowest density in the hindbrain with 154 ±19 cells per mm^2^ (mean ±SEM) to the highest density of ORF1p+ cells in the isocortex with 451 ±44 cells per mm^2^ (mean ±SEM) and the thalamus with 446 ±50 cells per mm^2^ (mean ±SEM). The proportion of ORF1p+ cells per anatomical region fluctuated between 10% ±2.1 (ventral striatum, mean ±SEM) and 31% ±1.6 (thalamus, mean ±SEM). The dorsal striatum (“striatal dorsal” in the Allen Brain Atlas denomination) exhibited the lowest ORF1p expression intensity (658 ±3 mean ±SEM) of all regions tested, the hindbrain the highest mean intensity of ORF1p per cell (mean ±SD 1221 ±548) as illustrated in Figure 1B and quantified in Figure 1D and 1F. Interestingly, cell density did not correlate with expression levels. Dorsal and ventral striatum for instance displayed similar ORF1p intensities per cell but exhibited significant differences in ORF1p cell density and proportion. The “midbrain motor” region as defined by the Allen Brain Atlas showed an intermediate cell density (mean ±SEM 265 ±16 cells per mm^2^) and a rather high ORF1p expression intensity (mean ±SEM 1006 ±533). Statistical analysis comparing mean density of ORF1p+cells per mm^2^ or mean intensity per ORF1p+ cells among regions confirmed the heterogeneity concerning ORF1p expression throughout the mouse brain (Fig. 1D, 1F). Slide scanner and confocal images revealed an exceptionally high ORF1p expression intensity in the ventral region of the midbrain which we identified as the *Substantia nigra pars compacta (SNpc)*. This region displayed an important density of ORF1p+ cells and a comparatively high level of ORF1p expression as illustrated by confocal imaging (Fig. 1B, panel 8), but could not be quantified independently with our brain-wide approach due to the geometrically-complex anatomy of this region and its small size (subregion-level in the Allen Brain Atlas hierarchy). Another region which could not be included in our brain-wide analysis was the cerebellum due to its extremely high density of cell nuclei. However, slide scanner and confocal imaging (Fig. 1B, panel 10) revealed that ORF1p is expressed in Purkinje cells, while not detectable in the molecular or granular layers.

**Fig. 1:**
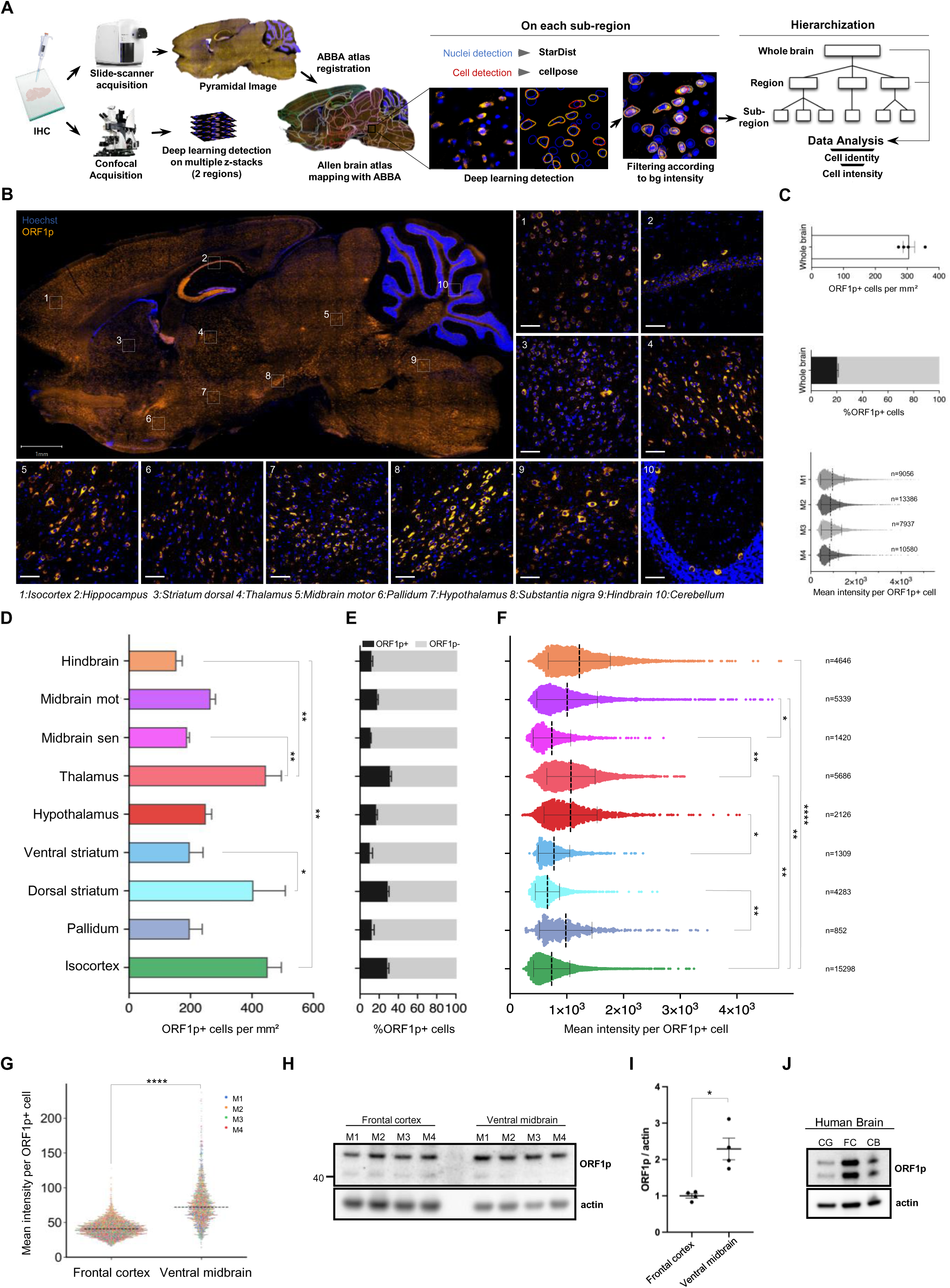
Widespread and heterogenous expression of ORF1p protein in the mouse brain. (A) Schematic representation of the unbiased cell detection pipeline on large-scale and confocal images. Immunofluorescent images on sagittal mouse brain slices were acquired on a digital pathology slide scanner or on a confocal microscope (DNA stain: Hoechst, neuronal marker; NeuN, protein of interest: ORF1p). Pyramidal images aquired with the slide scanner were then aligned with the hierarchical anatomical annotation of the Allen Brain Atlas using ABBA. Once the regions defined, a deep-learning based detection of cell nuclei (Hoechst staining, Stardist) and cell cytoplasm (NeuN staining, Cellpose) was performed on each sub-region of the atlas. Objects were filtered according to the background intensity measured in each sub-region for each channel (NeuN and ORF1p). The identity and intensity measures were analyzed at the regional and whole brain level. In parallel, confocal images (multiple z-stacks) of two selected regions (frontal cortex and ventral midbrain) were also acquired and identity and intensity were quantified using Cellpose. (B) Widespread and heterogenous expression of the LINE-1 encoded protein ORF1p in the mouse brain. Representative image of ORF1p immunostaining (orange) of a sagittal section of the brain of a young (three months-old) mouse acquired on a slide scanner. Scale bar = 1mm. (1-10) Representative images of confocal images of immunostainings showing ORF1p expression (orange) in 10 different regions of the mouse brain. Nuclei are represented in blue (Hoechst), scale bar = 50µm. (1) Isocortex, (2) Hippocampus, (3) Striatum dorsal, (4) Thalamus, (5) Midbrain motor, (6) Pallidum, (7) Hypothalamus, (8) Substantia nigra pars compacta, (9) Hindbrain, (10) Cerebellum. ORF1p expression profile in the mouse brain. The entire mouse brain with the exception of the olfactory bulb and the cerebellum were analyzed according to the pipeline on large-scale images described in (A). (C) Bar plot showing the total number of ORF1p+ cells per mm² in the mouse brain. Data is represented as mean ±SEM, n=4 mice (top). Bar plot indicating the proportion of ORF1p+ cells compared to all cells detected. Data is represented as mean ±SEM, n=4 mice (labeled M1 to M4), 202001 total cells analyzed (middle). Scatter plot showing the mean intensity of ORF1p per ORF1p+ cell. Data is represented as mean ±SD, n=4 mice, 40999 ORF1p+ cells analyzed (bottom). (D-F) ORF1p expression profile (density, proportion and expression) in defined anatomical regions of the mouse brain. Nine anatomical regions as defined by the Allen Brain Atlas and mapped onto sagittal brain slices (four three-month-old Swiss/ OF1) with ABBA were analyzed using the pipeline on large-scale images described in (A). (D) ORF1p+ cell density in 9 different regions. Bar plot showing the number of ORF1p+ cells per mm². Data is represented as mean ±SEM; *p<0.05; **p<0.01; adjusted p-value, one-way ANOVA followed by a Benjamin-Hochberg test (E) Proportion of ORF1p positive cells in 9 different regions. Bar plot showing the proportion of ORF1p+ cells among all cells detected per region. Data is represented as mean ±SEM. (F) Mean ORF1p expression per cell in 9 different regions. Dot plot showing the mean intensity of ORF1p signal per ORF1p+ cell in 9 different regions. Data is represented as mean ±SD. The number of analyzed cells per region is indicated in the figure. *p<0.05; **p<0.01; ***p<0.001; ****p<0.0001; adjusted p-value, nested one-way ANOVA followed by Sidak’ multiple comparison test. (G) ORF1p expression in the mouse frontal cortex and ventral midbrain. Confocal images with multiple z-stacks. Dot plot representing the mean intensity levels of ORF1p per ORF1p+ cells. Four three-month-old Swiss/ OF1 mice (labeled as M1 to M4) are represented each by a different color, the scattered line represents the median. ****p<0.0001, nested one-way ANOVA. Total cells analyzed = 4645. (H-I) ORF1p expression in the mouse frontal cortex and the ventral midbrain. (H) Western blots showing ORF1p (top) and actin expression (bottom) in four individual mice per region which were quantified in (I) using actin as a reference control. The signal intensity is plotted as the fold change of ORF1p expression in the ventral midbrain to ORF1p expression in the frontal cortex. *p<0.05; two-sided, unpaired student’s-test. (J) ORF1p expression in three regions of the human brain. Western blot showing human ORF1p expression in the cingulate gyrus (CG), frontal cortex (FC) and cerebellum (CB) of post-mortem tissues from a healthy individual. ORF1p (Top), Actin (bottom). For the full Western blot image please see Suppl Fig. 2A.

In order to confirm ORF1p expression by an independent method, we performed Western blot analysis on six micro-dissected regions from the mouse brain (Swiss/OF1 mouse, three-month old). As shown in Suppl Fig. 1E, ORF1p is expressed in all six regions with varying expression levels confirming the overall presence of ORF1p throughout the brain. We then chose two regions with significantly divergent ORF1p expression intensities as identified and quantified on pyramidal large-scale images: the frontal cortex (low) and the ventral midbrain (intermediate to high). We confirmed a significant higher expression of ORF1p in the ventral midbrain compared to the frontal cortex using an approach based on the unbiased, automated quantification of multiple z-stacks using a confocal microscope (Fig. 1G) and by Western-blotting on micro-dissected regions (Fig. 1H, 1I). In concordance with the findings stemming from the large-scale image quantification pipeline (Fig. 1F), the ventral midbrain showed ≈ 2-times higher expression of ORF1p than the frontal cortex as quantified in Figure 1G (1.8-fold) and Figure 1I (2.3-fold) validating our cellular detection methodology for pyramidal large-scale imaging and underscoring the heterogeneity of ORF1p expression levels in the mouse brain.

To investigate intra-individual expression patterns of ORF1p in the post-mortem human brain, we analyzed three brain regions of a neurologically-healthy individual (Fig. 1J, entire Western blot membrane in Suppl Fig. 2A) by Western blotting using a commercial and well characterized antibody which we further validated by several means. While there is some discrepancy in the field, the double band pattern in Western blots has been observed in other studies for human ORF1p outside of the brain ^40, 41^ as well as for mouse ORF1p ^42^. The nature of this lower band is unknown, but it might be due to truncation ^43^, specific proteolysis or degradation. Nevertheless, it appears that in cell culture models, a single ORF1p band is observed, whereas in murine and human samples, the ORF1p band is, to our knowledge, consistently associated with a lower molecular weight band ^35–37, 40–42, 44, 45^. We validated the antibody by immunoprecipitation and siRNA knock-down in human dopaminergic neurons in culture (differentiated LUHMES cells, Suppl Fig. 2B and 2C) where we detected in most cases the upper band only. ORF1p was expressed at different levels in the human post-mortem cingulate gyrus, the frontal cortex and the cerebellum underscoring a widespread expression of human ORF1p across the human brain. This was in accordance with ORF1p immunostainings of the human post mortem cingulate gyrus (Fig. 2H and Suppl Fig. 2E) and frontal cortex (Suppl Fig. 2E), with an absence of ORF1p staining when using the secondary antibody only (Suppl Fig. 2E).

**Fig. 2:**
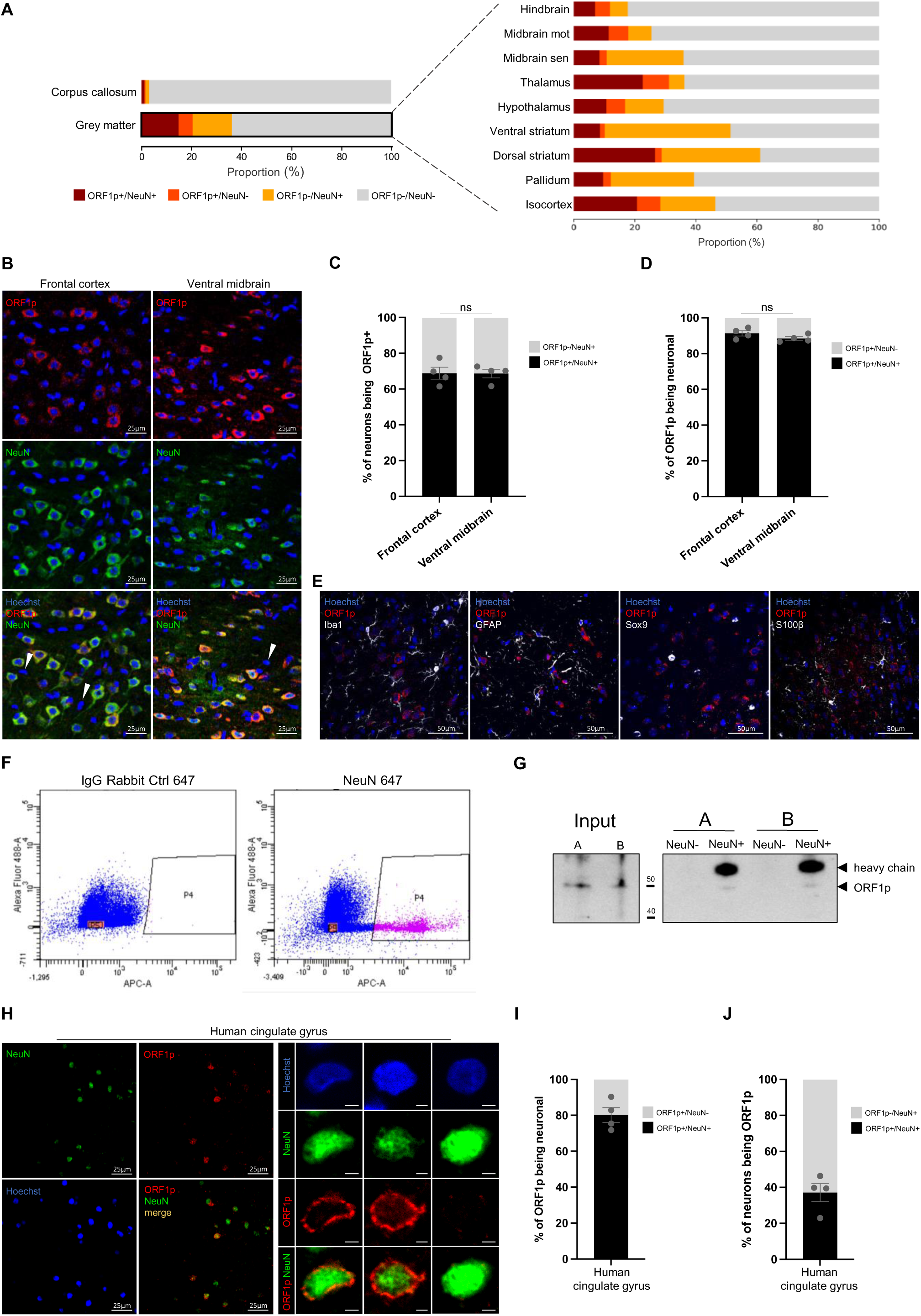
ORF1p is predominantly expressed in neurons in the mouse brain. (A) ORF1p expression is absent in the white matter (corpus callosum) and predominantly expressed in neurons. Proportion of ORF1p+/NeuN+, ORF1p+/NeuN-, ORF1p-/NeuN+ and ORF1p-/NeuN- cells in the white matter (corpus callosum) and the grey matter (left) and in nine different regions (right) analyzed by the cell detection pipeline on large-scale images presented in Figure1A. Exact values can be found in Suppl_Table1. (B) Representative confocal microscopy images showing ORF1p (red) and NeuN expression (green) in two different regions of the mouse brain. The bottom images show the merge of the two stainings, an overlap of both markers is represented in orange. z-projections; scalebar = 25µm. (C) Proportion of neurons expressing ORF1p in the frontal cortex and ventral midbrain quantified on confocal images. ns: non-significant; chi-square test on the cell number of the different cell-types analyzed; n=4 mice, data is represented as mean ± SEM. (D) Proportion of ORF1p+ cells identified as NeuN+ or NeuN- in two different regions, analyzed by confocal microscopy on multiple z-stacks. ns: non-significant; chi-square test, n=4 mice. (E) ORF1p does not colocalize with glial or microglial cell markers. Representative confocal microscopy images showing ORF1p staining (red) and three different glial cell (GFAP, Sox9, S100β) or microglial (Iba1) markers (white). Note that Iba1 antibody (rabbit) was used with the ORF1p 09 antibody (guinea pig, in house) z-projections, scalebar = 50µm. (F-G) Separation of neuronal and non–neuronal - cells by FACS confirms predominant neuronal expression of ORF1p. (F) Neuronal (NeuN+) and non-neuronal (NeuN-) cells isolated by fluorescent activated cell sorting (FACS). Dot plots showing autofluorescence versus an appropriate control antibody (IgG rabbit 647; left) and an antibody against NeuN (AB 657, right). The P4 window represents isolated NeuN+ cells (pink) and the P5 fraction NeuN- cells (orange) containing the same number of cells as sorted in P4 for comparison, others NeuN- are represented in blue. (G) Western blot showing ORF1p expression in NeuN- and NeuN+ FACS- sorted cells stemming from Figure F (A and B representing two different FACS experiments). (H) Representative confocal microscopy images showing ORF1p (red), NeuN (green) and Hoechst (blue) in the cingulate gyrus of the human brain. z-projection; scalebar = 25µm (left). Examples of individuals neurons expressing ORF1p or not are shown on the right panel. z-projection; scalebar = 5 µm (right). (I) Proportion of ORF1p+ cells identified as NeuN+ or NeuN- in the human cingulate gyrus, analyzed by confocal microscopy on multiple z-stacks. (J) Proportion of neurons expressing ORF1p in the human cingulate gyrus, analyzed by confocal microscopy on multiple z-stacks.

In summary, our findings reveal the consistent presence of ORF1p expression throughout the mouse brain in all anatomical regions analyzed with high regional variability in terms of density of ORF1p+ cells and ORF1p+ cell intensity. ORF1p is also expressed in the human brain in at least three brain regions. This finding raises several questions concerning cell-type identity of ORF1p expressing cells and potential functions or consequences of ORF1p expression in the mouse and human brain at steady-state.

### ORF1p is predominantly expressed in neurons

Following our observation of a wide-spread expression of endogenous ORF1p throughout the brain, we first addressed the question of the cellular identity of ORF1p+ cells. To this end, we used the neuron-specific marker NeuN, commonly used to identify post-mitotic neurons in the central nervous system ^46^. This allowed us to determine the proportion of neuronal (NeuN+) or non-neuronal cells (NeuN-) expressing ORF1p (ORF1p+) or not (ORF1p-). Making use of our large-scale imaging approach (Fig. 1A), we observed drastic dissimilarities in detected cellular proportions between the white and grey brain matter. As expected, we observed only 1% of NeuN+ cells in the white matter (corpus callosum; Fig. 2A) validating both, the neuronal marker NeuN as such and the ABBA superposition of the Allen Brain Atlas onto the sagittal brain slices. In the grey matter, our approach detected 30.5% NeuN+ cells (dark red and yellow bars in Fig. 2A) which, according to the literature, should include all post-mitotic neurons with only minor exceptions ^46–49^ and corresponds to the reported proportion of neurons present in the mouse brain ^50^. The nine identified grey matter regions in Fig. 2A display the proportions of the different cell types per region. The proportion of all cells in a given region which are positive for ORF1p (dark red bars) differed between regions (lowest proportion: hindbrain: 7%; highest proportion dorsal striatum: 26.6%). In the isocortex and the midbrain motor-related regions, the majority of neurons detected express ORF1p (54% and 59% by large-scale analysis, Suppl Fig. 3A; 68.7% and 68.8% by confocal imaging, Fig. 2B, quantified in C), while in the midbrain sensory related regions the proportion dropped to 25% whereas it reached 82% in the thalamus (Suppl Fig. 3A). Altogether, nearly half of all NeuN+ cells throughout the mouse brain expressed ORF1p (mean of all regions: 48.2%; Suppl Fig. 3A). Regarding the cell identity of ORF1p+ cells brain-wide, more than 70% were identified as neuronal by the large-scale approach (Suppl Fig. 3B). This contrasted somewhat with results obtained by the second approach using confocal imaging on multiple z-stacks which indicated that 91.3% (frontal cortex) and 88.5% (ventral midbrain) of ORF1p+ cells were neuronal (Fig. 2D). This difference in percentages of ORF1p+ expressing neurons among all neurons between the large-scale image cell detection methodology and the confocal workflow is most probably due to technical limitations inherent to the large-scale pipeline. Indeed, with the latter approach, region-dependent differences in cell density and signal intensity might be the cause for an underestimation of the proportion of ORF1p+ cells being neuronal due to difficulties in cell detection by StarDist/Cellpose (high cell density) on a single focal plan, technical difficulties which are widely reduced by the multiple z-stack based approach when using a confocal microscope. Moreover, the large-scale pipeline involves background measurements in each sub-region in order to apply stringent filtering. This, however, results in a loss of true positives cells, but avoids cells which are out-of-focus, the presence of which is inherent to the slide scanner microscope which lacks optical sectioning (see Materials & methods section for evaluated model performance). Notably, frontal cortex and ventral midbrain present similar proportion of neurons expressing ORF1p (Fig. 2C), although the percentage of NeuN+ cells between these two regions is significantly different (Suppl Fig. 3C). As we could not rule out that ORF1p might also be expressed in non-neuronal cells, we turned to non-neuronal markers specific for different glial cell populations using two different astrocytic markers (GFAP, Sox9), the astro-and oligodendrocytic marker S100β and the microglial marker Iba1 ^50, 51^ and performed co-staining with ORF1p followed by confocal imaging as illustrated in Figure 2E. We screened multiple images of frontal cortex, ventral midbrain, hippocampus and striatum and did not find a single ORF1p+ cell which could unambiguously be defined as non-neuronal. This indicated that ORF1p is not or only very rarely expressed in non-neuronal cells. To further confirm the predominant presence of expression of ORF1p in neurons and the absence of ORF1p expression in non-neuronal cells, we used fluorescence-activated cell sorting (FACS) to isolate neurons (using a NeuN antibody) and non-neuronal cells (NeuN-) from the adult mouse brain followed by Western blotting with an antibody against ORF1p (Fig. 2F, 2G). As described above, this antibody is well characterized, extensively used and was validated further in this study. After FACS-sorting of neurons from the adult mouse brain using an antibody against NeuN (Fig. 2F, G), we detected ORF1p exclusively in the neuronal population (NeuN+, internal control = heavy chain), confirming the results based on two different imaging approaches. Finally, to assess whether predominant, if not exclusive ORF1p expression in neurons is mouse brain specific or a pattern also applicable to the human brain, we investigated the identity of ORF1p expressing cells in the post-mortem cingulate gyrus of a healthy human brain. Similar to what we found in the mouse brain, we observed sparse NeuN expression in the white and extensive NeuN staining in the grey matter corresponding to the cortical layers (Suppl Fig. 2D, grey and white matter separated by a dashed line) with ORF1p+ cells predominantly located in the grey matter (confocal images in Fig. 2H, Suppl Fig 2E are located in the grey matter). All cells stained by ORF1p were co-stained with NeuN indicating that ORF1p was expressed in neuronal cells in the human brain (Fig. 2H). However, due to the lower signal quality inherent to human post-mortem sections compared to mouse sections, the identity of ORF1p+ cells was estimated to be 80% neuronal by the automated image analysis pipeline of confocal images (Fig. 2I), although no ORF1p+ / NeuN- cells could be clearly identified. Of all neurons identified, 37.2% were ORF1p+ (Fig. 2J), indicating that, similar to the mouse brain, only a fraction of neurons express ORF1p (Fig. 2H, right).

Next, we asked the question of a neuron-subtype specific expression of ORF1p. Our previous study had revealed a higher expression of ORF1p in tyrosine-hydroxylase (TH) positive neurons compared to TH-negative neurons in the mouse ventral midbrain ^17^. A recent study reported that endogenous LINE-1 RNA and ORF1p expression were higher in parvalbumin (PV)-positive interneurons compared to PV-negative neurons in the mouse hippocampus ^18^. To address the question of a generalized co-expression of ORF1p and PV cells, we co-stained sagittal brain sections of young mice with antibodies against ORF1p and PV and, in some cases the lectin Wisteria floribunda agglutinin (WFA), which specifically stains glycoproteins surrounding PV+ neurons. Confocal imaging on several brain regions including the hippocampus, cortex, cerebellum, hindbrain, ventral midbrain and thalamus revealed ORF1p+ neurons co-expressing PV, but also many examples of equally intense ORF1p+ neurons that do not express PV (Suppl Fig. 4).

In summary, ORF1p expression in the mouse and human brain is widely restricted to neurons of which a proportion express ORF1p. This raises the question of the function and consequences of ORF1p expression specifically in neurons but also on the dynamic regulation of this expression upon exogenous (exposome) or endogenous (aging) challenges.

### ORF1p expression is increased in the aged mouse brain

ORF1p is expressed at steady-state throughout the brain, but whether this expression is dynamically regulated remains unknown. Aging has been linked to LINE-1 regulation in some studies ^16, 52^ potentially as both, a trigger and as a consequence of LINE-1 activation, but whether this is true for the brain and if yes, whether this might be region-specific has not been investigated brain-wide. We therefore addressed the question whether advanced age was paralleled by a change of expression patterns or expression levels of ORF1p in the brain. We first analyzed ORF1p expression levels comparing young (3-month) to aged (16-month) mouse brains using the cell detection workflow applied to large-scale images described in Figure 1A. Interestingly, the mean intensity of ORF1p expression increased moderately but significantly with advanced age throughout the brain (13% increase brain-wide; n=4 young mice; n=4 aged mice; p=0.03; Fig. 3A). This was in contrast to another protein, NeuN, which we used as a control and whose intensity did not change between young and aged brains (n=4 young mice; n=4 aged mice; p=0.27; Fig. 3B). Frequency distribution analysis unveiled a shift in ORF1p mean expression per cell in aged mice (Fig. 3C). Importantly, the Hoechst mean intensity within nuclei of ORF1p+ cells, serving as an internal control, showed no significant change (Fig. 3D). Among nine analyzed regions, five demonstrated a general increase in ORF1p mean intensity per cell in aged mice (p≤0.05), a change independent from inter-individual variations in both young and aged mice (Fig. 3E). The increase of ORF1p expression (fold change intensity) throughout the brain, reaching nearly a 30% increase in some regions, is represented on the heatmap in Figure 1F. These results were confirmed by the confocal imaging approach; ORF1p expression in the frontal cortex remained unchanged but increased significantly in the ventral midbrain region in aged mice as shown in Figure 3G and quantified in Figure 3H.

**Fig. 3:**
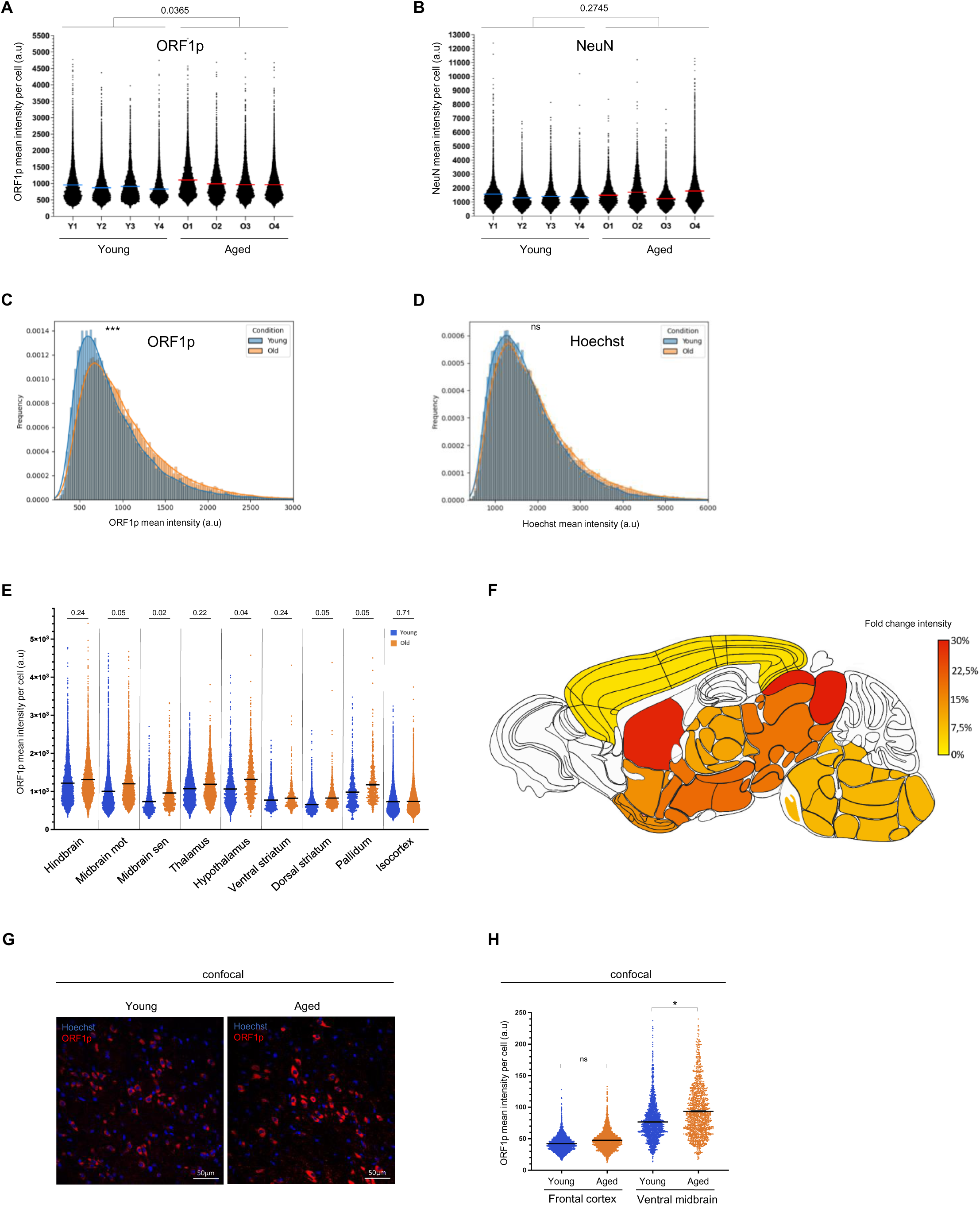
ORF1p expression is increased in some regions of the aged mouse brain. (A) ORF1p mean intensity per ORF1p+ cell in the brain, analyzed on large-scale images. Dot plot showing the ORF1p mean expression per ORF1p+ cell in young (Y1-4) and aged (O1-4) mice in the whole brain (except cerebellum and olfactory bulb). 74985 total cells were analyzed; * p<0.05, nested two-way ANOVA; n=4 mice per condition, data is represented as mean ± SEM. (B) NeuN mean intensity per NeuN+ cell in the brain, analyzed on large-scale images. Dot plot showing the NeuN mean expression per NeuN+ cell in young (Y1-4) and aged (O1-4) mice in the whole brain (except cerebellum and olfactory bulb). Nested two-way ANOVA; n=4 mice per condition, data is represented as mean ± SEM. (C) Frequency distribution of ORF1p mean intensity in ORF1p+ cells of young (blue) and aged (orange) mice. ***p<0.001, Kolmogorov-Smirnov test. (D) Frequency distribution of Hoechst mean intensity in the nuclei of OrF1p+ cells of young (blue) and aged (orange) mice. ns: non-significant, Kolmogorov-Smirnov test. (E) Mean ORF1p expression per ORF1p+ cell in nine different anatomical regions. Dot plot showing the ORF1p mean expression per ORF1p positive cell (n=74985). Adjusted p-value are represented, two-tailed nested t-test followed by a Benjamin, Krieger and Yukutieli test; n=4 mice per region, data is represented as mean ± SEM. (F) Color-coded representation of fold-changes of ORF1p expression comparing young and aged brains. Represented is the fold-change in percent (aged vs young) of the “mean of the mean” ORF1p expression per ORF1p+ cell quantified and mapped onto the nine different regions analyzed as shown in (J). (G) Representative confocal microscopy images showing increased ORF1p expression (red) in the ventral midbrain region of aged mice (one z plan is shown). Cell nuclei are shown in blue (Hoechst staining). Scalebar = 50µm. (H) ORF1p expression is increased in the ventral midbrain of aged mice. Dot plot representing ORF1p expression in two different regions of young and aged mice analyzed on confocal images with multiple z-stacks; total cells analyzed = 8381 ns: non-significant *p<0.05, two-tailed one-way ANOVA; dashed lines represent the medians.

We then asked whether the increase in ORF1p expression levels observed in several brain areas in aged compared to young mice was also accompanied by a change in expression patterns. We therefore analyzed cell proportions and densities comparing young and aged mouse brains. Globally, we observed a reduction in the proportion of ORF1p+/NeuN+ cells in aged mouse brains using the cell detection workflow applied to large-scale images described in Figure 1A, a phenomenon mainly driven by the midbrain motor, the dorsal striatum, the pallidum and the thalamus regions (Suppl Fig. 5A, dark red bars, Suppl_Table1). The confocal approach applied to two regions, the frontal cortex and the ventral midbrain (Suppl Fig. 5B), confirmed this reduction in ORF1p+/NeuN+ cell proportions in favor of the ORF1-/NeuN- cell population in the ventral midbrain with no change in cell proportions in the frontal cortex in accordance with the large-scale imaging approach (Suppl Fig. 5A). The predominantly neuronal identity of ORF1p+ cells, however, was unchanged in the ventral midbrain (Suppl Fig. 5C) just as the proportion of neurons expressing ORF1p (Suppl Fig. 5D). We observed a significant shift in NeuN+/- cell proportions (Suppl Fig. 5E) which could either suggest a decrease in NeuN+ cells or a gain of NeuN- cells in this region with age. While proportions are less sensitive to technical variability and can identify cell population shifts, cell densities allow for absolute comparisons. When quantifying global cellular densities throughout the brain, we did not observe a significant reduction of ORF1p expressing cells (Suppl Fig. 5F), neurons (Suppl Fig. 5G) or non-neuronal cells (Suppl Fig. 5H). When analyzing cell densities in the nine brain regions separately, there were no significant changes in ORF1p positive (Suppl Fig. 5I), NeuN positive (Suppl Fig. 5J) or NeuN negative (Suppl Fig. 5K) cell densities in eight out of nine brain regions. The only exception was the dorsal striatum, but technical limitations applying to this particular brain region might account for these changes. Indeed, the dorsal striatum is different from the other brain regions as it represents the only region consisting of a single ABBA sub-region resulting in only one overall background measurement. Taken together, while there were no major changes in cell proportions, densities nor in ORF1p+ cell identities, we observed an age-dependent increase in ORF1p expression per cell of up to 27%.

### Coding LINE-1 transcripts are increased in aged human dopaminergic neurons

Following the observation of increased ORF1p expression in the aged mouse brain, among which the ventral midbrain, and given the age-related susceptibility of dopaminergic neurons in the *SNpc* to cell death and to degeneration in PD ^53^, we turned to a previously published RNA-seq dataset of laser-captured micro-dissected post-mortem human dopaminergic neurons of brain-healthy individuals ^54^, in order to interrogate full-length LINE-1 mRNA expression profiles as a function of age. To avoid read-length bias to which TE analysis is particularly sensitive, we analyzed only the data derived from 50bp paired-end reads of linearly amplified total RNA as this dataset represented all age categories (n=41; with ages ranging from 38 to 97; mean age: 79.88 (SD ±12.07); n=6 ≤65y; n=35 >65y; mean PMI: 7.07 (SD ±7.84), mean RIN: 7.09 (±0.94), metadata available in Suppl_Table5). As age-related dysregulation of TEs might not be linear, we considered individuals with ages-at-death younger or equal to 65 years as “young” (n=6, 38-65 years, mean age 57.5 years (SD ±9.9)) and individuals older than 65 years as “aged” (n=35, 65-97 years, mean age 83 years (SD±7.8). The expression of the dopaminergic markers tyrosine hydroxylase (TH) and LMX1B were similar in both populations indicating no apparent change of dopaminergic identity of analyzed melanin-positive dopaminergic neurons (Suppl Fig. 6A, B). Next, we compared the expression of repeat elements at the class, family and name level based on the repeat masker annotation implemented in the UCSC genome browser using a commonly used mapping strategy for repeats consisting of randomly assigning multi-mapping reads ^55^. No overt dysregulation of repeat elements at either level of repeat element hierarchy was observed (Suppl Fig. 6C-F). There was a modest but significant increase in several younger LINE-1 elements including L1HS and L1PA2 at the “name” level (Fig. 4A, B), an analysis which was however underpowered (post-hoc power calculation; L1HS: 28.4%; L1PA2: 32.8%) and thus awaits further confirmation in independent studies. No expression changes were observed for HERVK-int, a human endogenous retrovirus family with some copies having retained coding potential (Fig. 4B) or other potentially active TEs like HERVH- int, HERV-Fc1, SVA-F or AluYa5 transcripts in the >65y group (Suppl Fig. 6G). Interestingly, L1HS expression was highly correlated with L1PA2 expression and this correlation extended to almost all younger LINE-1 subfamilies weaning down with evolutionary distance (Fig. 4C). This was not true for other active TEs as L1HS was negatively correlated with HERVK-int expression (Fig. 4C). Several regulators of LINE-1 activity have been identified ^17, 56^ and correlation of their expression with L1HS might allow to infer their relevance of interaction (activation or repression) with L1HS in human dopaminergic neurons. Spearman correlation analysis revealed three known repressors of LINE-1 activity whose expression was negatively correlated with LINE-1 expression; EN1 (Engrailed 1 ^17^, Suppl Fig. 7A) with important functions for dopaminergic neuron homeostasis ^57^, CBX5/HP1a, a heterochromatin binding protein binding to the histone mark H3K9me3, thereby mediating epigenetic repression ^58^ (Suppl Fig. 7B) and XRCC5/6, also known as Ku86/Ku70, which are essential for DNA double-stranded break repair through the nonhomologous end joining (NHEJ) pathway and limit LINE-1 full-length insertions ^59^ (Suppl Fig. 7C). The transcripts of these genes showed, although not statistically significant, a trend for decreased expression in the elderly (Suppl Fig. 7D-G). Based on the increase of young LINE-1 families L1HS and L1PA2 in aged human dopaminergic neurons and the finding that ORF1p was increased in the aged mouse brain, we focused our attention on LINE-1 elements with coding potential for ORF1 and ORF2 according to the L1Basev2 annotation which are specific elements comprised in the L1HS and L1PA2 annotation at the “name” level. Most of the 146 full-length and coding LINE-1 termed UIDs (= Unique Identifier) in the L1Base are L1HS elements (76.03%), whereas the remaining 35 UIDs belong to the evolutionary older L1PA2 family (Suppl Fig. 7A). The L1Base annotation is based on the human reference genome (GRCh38) and annotates 146 human full-length (>6kB), intact LINE-1 elements (ORF1 and ORF2 intact) with a unique identifier from 1 to 146 ^4^. Attribution of sequencing reads to a specific, individual TE copy is problematic ^60^ and several approaches have been proposed to circumvent this problem including the mapping of unique reads ^55^.While several tools using expectation maximization algorithms in assigning multi-mapping reads have been developed and successfully tested in simulations ^55, 61^, we used a different approach in mapping unique reads to the L1Base annotation of full-length LINE-1. Specific “hot” LINE-1 loci in a given cellular context have been identified ^3^, but usage of the L1Base annotation enabled an unbiased approach albeit ignoring polymorphic LINE-1 sequences. Unique read mapping strategies for repeat elements, especially young LINE-1 elements, will unavoidably underestimate LINE-1 locus-specific expression levels ^55^, but will be most accurate in assigning reads to a specific genomic location while allowing the comparison of two different conditions analyzed in parallel. Of the 146 full-length LINE-1 elements in the L1Base annotation, 111 were of the L1HS family and 35 belonged to the L1PA2 family (Suppl Fig. 8A). Assuming that expression of UIDs was correlated with mappability, we plotted a mappability count of each UID against its mean normalized read count expression of the six individuals ≤65y (Suppl Fig. 8B, C). Non-parametric Spearman correlation revealed no correlation between UID mappability and expression (Suppl Fig. 8C) indicating no apparent bias between the two parameters. However, individual UID dependency of mappability on expression cannot be excluded, especially for high expressing UIDs like UID-16 for example (Suppl Fig. 8B, C). Expression of LINE-1 at the locus-level has been attributed to artefacts not representing autonomous transcription including differential high intronic read counts ^62^, pervasive transcription or reads attributable to passive co-transcription with genes when the LINE-1 element is intronic ^63^. To evaluate the latter, we determined the number of intronic (46.58%) and intergenic UIDs (78/146; Suppl Fig. 8D) and identified the corresponding genes for intronic UIDs (Suppl Fig. 8E). Of the 146 UIDs, 140 passed the threshold of >3 reads in at least 6 individuals. Differential expression of UID between “young” and “aged” dopaminergic neurons revealed several significantly deregulated full-length LINE-1 loci (Fig. 5A). Paired analysis of the expression of all UIDs indicated a general increase (Fig. 5B), especially of low expressed UIDs. The comparative analysis of the sum expression of UIDs per individual comparing young (≤65y) with elderly human dopaminergic neurons, however, did not reach statistical significance (Fig. 5C). Several specific loci were dysregulated, in particular UID-68 (Fig. 5A and 5D), a L1HS element located on chromosome 7 (chr7: 141920659-141926712) in between two genes, OR9A4 (olfactory receptor family 9 subfamily A member 4) and CLEC5A (C-type lectin domain containing 5A; Fig. 5E and Suppl Fig.7E). This specific full-length LINE-1 element had a high mappability count of 16 (range of all UIDs: 1-30, mean 9.0 (SD ±6.05), Suppl Fig. 7C) and a post-hoc power analysis score of 96.6% (continuous endpoint, two independent samples, alpha 0.05). To rule out any influence of “hosting” gene transcription interference on measurable UID-68 expression differences, we performed Spearman correlation which did not indicate any correlation between OR9A4 (Fig. 5F) or CLEC5A (Fig. 5G) expression with UID-68. Together, this indicated that UID-68 might be a candidate for an age-dependent gain of activity. Other dysregulated UIDs (i.e. UID-129, UID-37, UID-127 and UID-137) had either a low mappability score, a low post-hoc power or did not pass the visualization check in IGV, reinforcing the notion that a combination of quality control criteria is crucial to retain a specific locus with confidence. In conclusion, TE expression analysis of this human dataset covering an age-span of 59 years (mean age difference between both groups 25.5 years) indicates an increase in the expression of young LINE-1 elements including those which have coding potential in elderly dopaminergic neurons, particularly a specific full-length LINE-1 element on chromosome 7 (UID-68). A slight net sum increase of UID transcripts/cell might be sufficient for the production of “above steady-state” levels of ORF1p and ORF2p. Other TEs with coding potential, namely members of the HERV family, were not increased. Further, correlation analyses suggest that L1HS expression might possibly be controlled by the homeoprotein EN1, a protein specifically expressed in dopaminergic neurons in the ventral midbrain ^57^, the heterochromatin binding protein HP1 and the DNA repair proteins XRCC5/6.

**Fig. 4:**
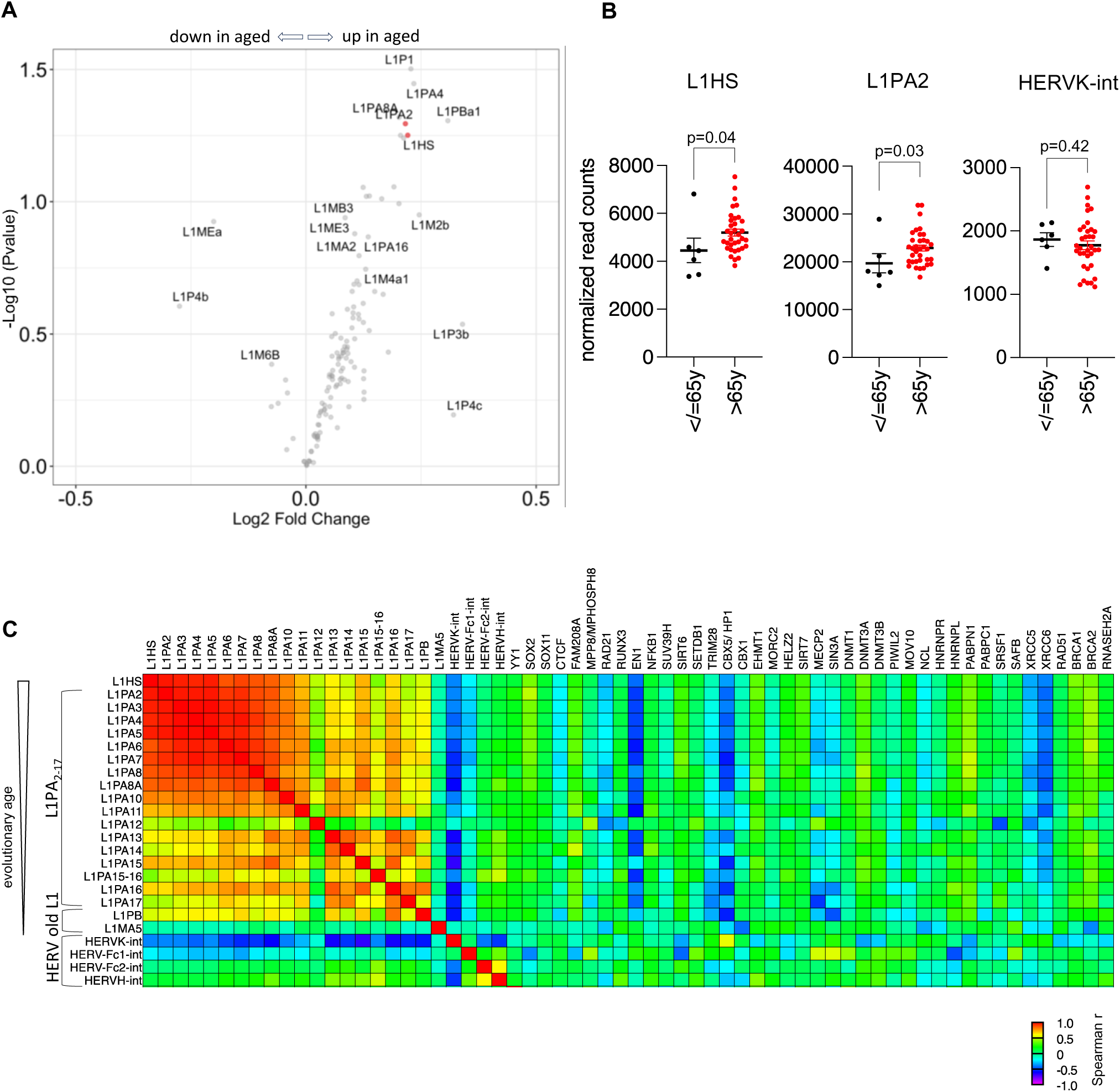
Young LINE-1 elements are increased in aged human dopaminergic neurons. TE transcript expression in RNA-seq data of laser-captured micro-dissected post-mortem human dopaminergic neurons of brain-healthy individuals was analyzed using RepeatMasker (multimappers) or the L1Base (unique reads). (A) Volcano plot of differential analysis of LINE-1 expression using DESeq2 comparing young (≤65y, n=6) or aged (>65y, n=35) human dopaminergic neurons at the “name” level of RepeatMasker. Young LINE-1 elements, including the two families L1HS and L1PA2 that have coding copies, are highlighted in red. (B) Scatter plots of normalized read counts (“name” level) of the young L1HS and L1PA2 families as well as the human endogenous virus family HERVK-int, another TE family with coding potential comparing young (≤65y, n=6) or aged (>65y, n=35) human dopaminergic neurons. Mann-Whitney test, p<0.05. (C) Correlation of the RNA expression levels of LINE-1 elements with known transposable element regulators in human dopaminergic neurons (all ages included). Spearman correlation of evolutionary close (L1HS, L1PA2-17) and distant LINE-1 (L1PB and L1MA5) as well as HERV elements with coding potential (HERV-Kint, HERV-Fc1, HERV-Fc2 and HERV-H-int) with known regulators of transposable elements for each individual sample, all ages included. HERV-W and TREX1 did not pass the normalized read count threshold of >3 reads in >6 individuals.

**Fig. 5:**
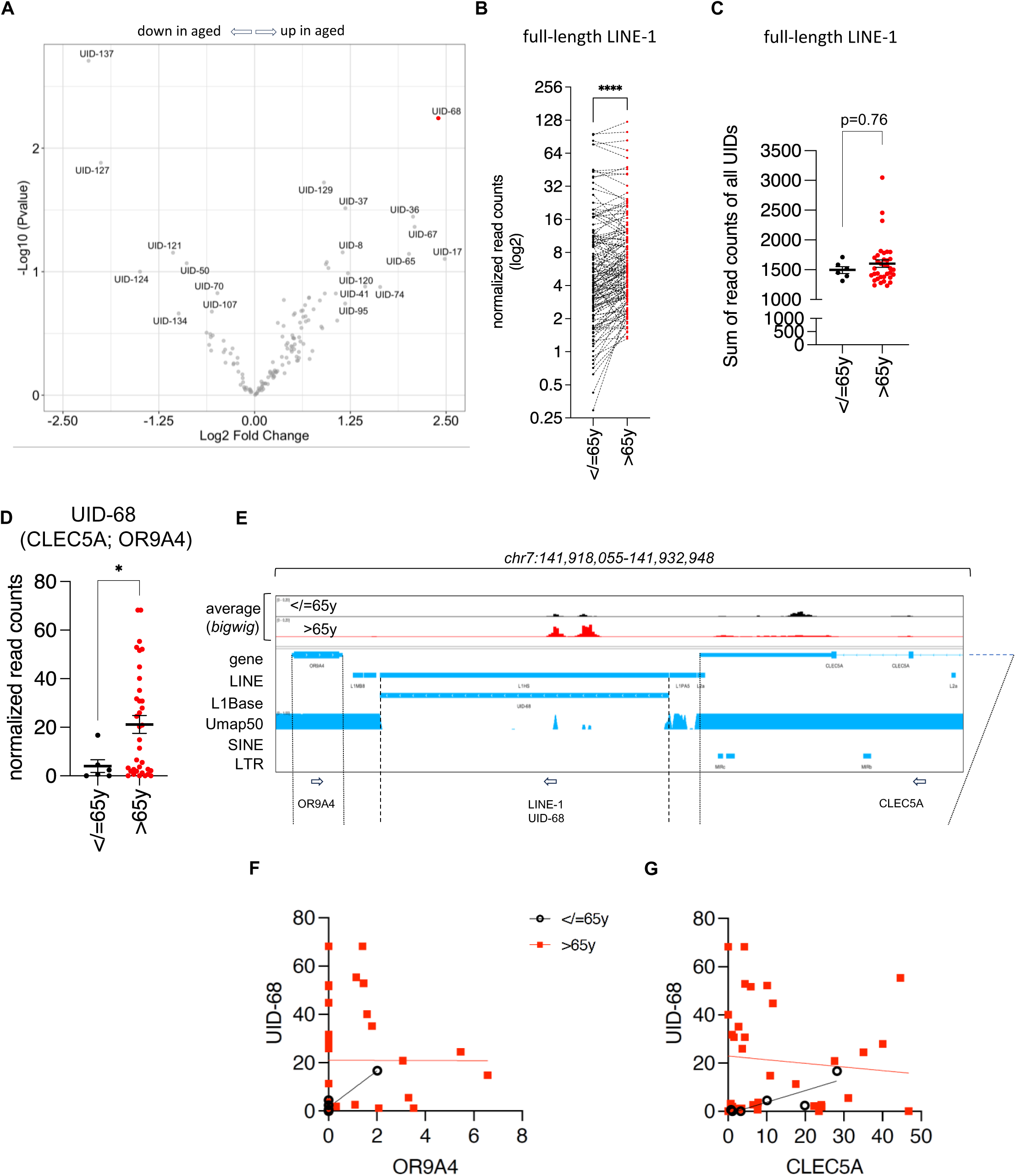
Dysregulation of locus-specific full-length LINE-1 elements in aged human dopaminergic neurons. (A) Volcano plot of differential expression analysis of TE expression using DEseq2 comparing young (≤65y, n=6) and aged (>65y, n=35) human dopaminergic neurons at the locus-level of specific full-length LINE-1 elements (140 of 146 “UID’s” as annotated in L1Base; threshold >3 reads in at least 6 individuals). (B) Pairwise comparison of the expression of 140 out of 146 full-length LINE-1 elements comparing young (≤65y, n=6) and aged (>65y, n=35) human dopaminergic neurons. Wilcoxon matched signed rank test, p<0.0001left panel). (C) The sum of read counts of all UIDs per individual were plotted comparing young (≤65y, n=6) and aged (>65y, n=35) human dopaminergic neurons. (D) UID-68 is dysregulated in aged human dopaminergic neurons. Normalized read counts of uniquely mapping reads mapping to the full-length LINE-1 element “UID-68” per individual were plotted comparing human post-mortem dopaminergic neurons from young (≤65y, n=6) and aged (>65y, n=35) individuals; two-tailed Mann-Whitney test (* p=0.046). (E) IGV window of the locus around the full-length LINE-1 UID-68 (chr7:141.918.055-141.932.948). UID-68 is located adjacent to the genes CLEC5A (right) and OR9A4 (left). Coverage and mappability of the locus including UID-68 is shown in tracks. Coverage is represented by average bigwig profiles (same scale, black: </=65y, red >65y); mappability of the genomic locus is depicted by Umap 50 tracks showing peaks overlapping with the peaks of reads (bigwig averages). (F) Spearman correlation analysis of the expression of UID-68 and OR9A4 in young (≤65y, n=6, black dots; Spearman r= 0.66, p=0.17)) or aged (>65y, n=35, red squares, Spearman r= 0.15, p=0.40) human dopaminergic neurons. (G) Spearman correlation analysis of the expression of UID-68 and CLEC5A in young (≤65y, n=6, black dots, Spearman r= 0.75, p=0.11) or aged (>65y, n=35, red squares; Spearman r= - 0.03, p=0.87) human dopaminergic neurons.

**Fig. 6:**
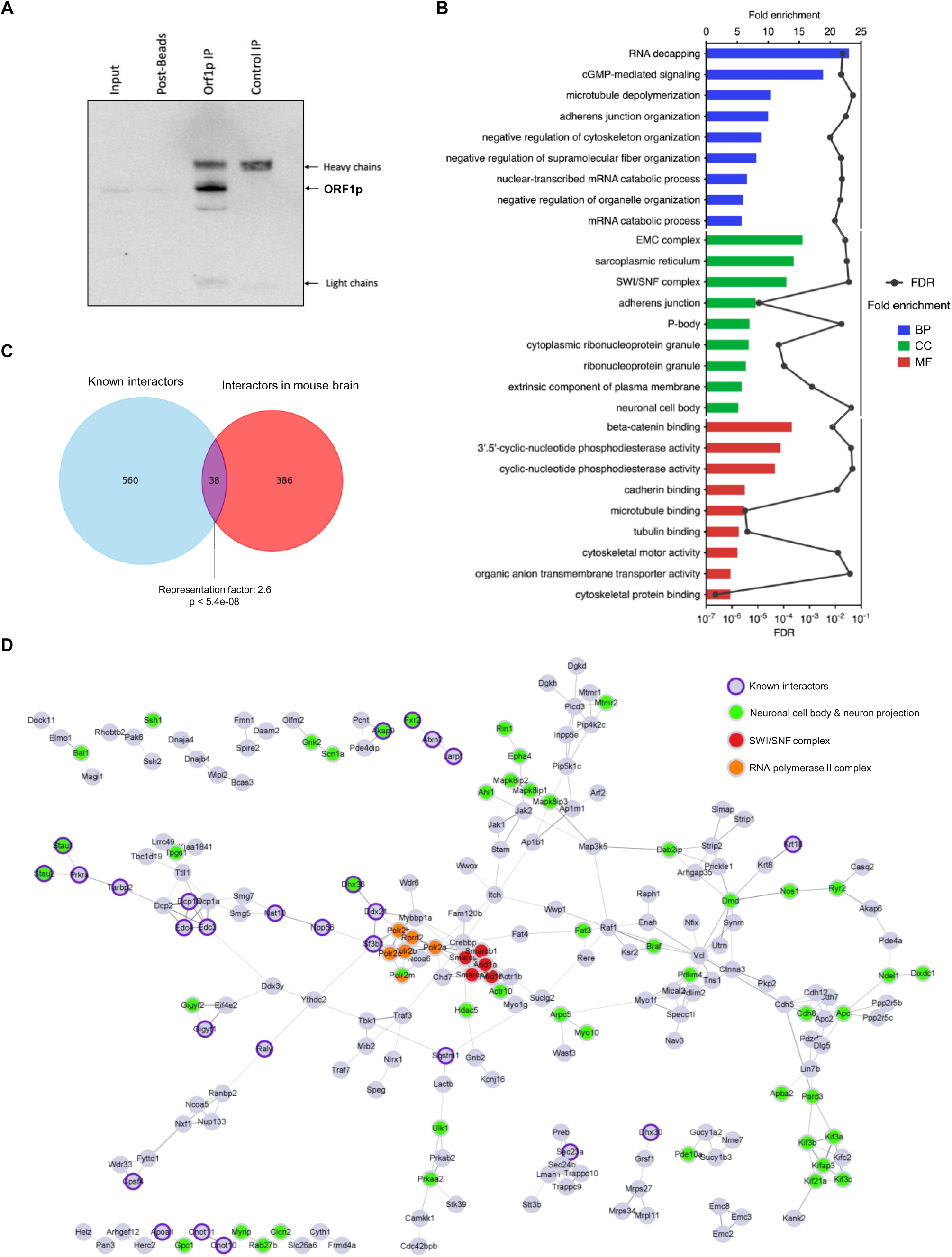
Endogenous ORF1p interactors in the mouse brain. Immunoprecipitation (IP) of endogenous ORF1p from the mouse brain. WB against ORF1p showing ORF1p enrichment after IP but no signal in the IgG control. Five independent samples were then prepared for proteomic analysis by mass spectrometry (LC-MS/MS). (B) GO slim enrichment analysis of proteins selected as endogenous ORF1p protein partners in the mouse brain after quantitative LC-MS/MS. ORF1p-immunoprecipitated proteins were categorized into GO slim terms. The nine GO slim term with the highest fold-change are plotted. Fold enrichment is depicted on the upper axis and displayed as bars, the FDR value appears on the lower axis and is represented by the black points. BP: Biological Process, CC: Cellular Component, MF: Molecular Function. (C) Venn diagram showing common interactors (purple) between interactors of endogenous ORF1p in the mouse brain identified in this study (red) and known (published) interactors of ORF1p (blue). Statistical significance of the overlap between the two groups of proteins was tested by an overrepresentation test (http://nemates.org/MA/progs/overlap_stats.html). (D) ORF1p associates with the SWI/SNF complex (red), RNA pol II complex (orange) and interactors belonging to GO terms related to neuronal cell body & neuron projection (green). Known interactors previously published ^35, 67, 72–76^ are indicated with a purple ring. STRING network of physical interactions where nodes represent proteins partners identified in (A) and edges thickness represents the strength of shared physical complexes. Only proteins sharing physical interactions were represented.

### Endogenous ORF1p interactors in the mouse brain

To go further in our understanding of steady-state neuronal ORF1p expression across the mouse brain, we immunoprecipitated ORF1p and performed quantitative label-free LC-MS/MS to identify potential protein partners of ORF1p in the adult mouse brain. We successfully immunoprecipitated endogenous ORF1p from whole brain lysates (Fig. 6A), where we detect ORF1p exclusively in the 5 independent ORF1p-IP samples (and not at all in 5 independent IgG-IP control samples; Suppl_Table2) and identified a total of 424 potential protein interactors associated with ORF1p (Suppl_Table2; n=5 mice). Using Gene Ontology (GO) analysis, we identified several interacting proteins belonging to GO terms related to known functions of the ORF1p protein in RNA binding, preferentially ^64^ but not exclusively *in cis* ^65^, for instance RNA decapping and mRNA catabolic process, or related to the known presence of ORF1p in ribonucleoprotein particles ^66, 67^ (GO: cytoplasmic ribonucleotide granule) or the presence of ORF1p in p-bodies ^65^ as shown in Figure 6B and listed in Suppl_Table3. Other GO terms that emerged, to our knowledge not previously associated with ORF1p, were related to cGMP-mediated signaling (GO: cGMP-mediated signaling and 3’-5’phosphodiesterase activity: i.e. PDE4A, PDE4B, PDE4DIP) and the cytoskeleton (GO: microtubule depolymerization, cytoskeleton organization, microtubule and tubulin binding, cytoskeletal motor activity and protein binding). cGMP signaling is regulated by 3’-5’ phosphodiesterases (PDEs) which degrade 3’,5’-cyclic guanosine monophosphate (cGMP) and 3’,5’-cyclic adenosine monophosphate (cAMP), an activity essential for cell physiology for the integration of extra- and intracellular signals including neuronal excitability, synaptic transmission and neuroplasticity ^68, 69^. Further, several ORF1p interacting proteins were constituents of the mating-type switching/sucrose nonfermenting complex (SWI/SNF complex), i.e. ARID1A, ARID1B, SMARCA2, SMARCB1, SMARCC2), an ATP-dependent chromatin remodeler complex disrupting nucleosome/DNA contacts to facilitate DNA/chromatin accessibility by shifting, removing or exchanging nucleosomes along DNA ^70, 71^. Finally, we also observed proteins belonging to the GO term “neuronal cell body”, corroborating with the neuron-specific presence of ORF1p in the brain. A comparative analysis with previous mass spectrometry studies ^35, 67, 72–76^ aimed at identifying ORF1p interacting proteins unveiled significantly more common proteins than randomly expected (overrepresentation test; representation factor 2.6, p< 5.4e-08; Fig. 6C), including LARP1, STAU2, ATXN2, RALY, TARBP2 or DDX21 (for a full list see Suppl_Table4). The presence of a significant number of overlapping ORF1p interactors in different non-neuronal human cells (HEK ^67, 72, 73^, HeLa ^74^, human breast and ovarian tumors ^76^ and hESCs ^75^) and mouse brain cells (our study), suggest conserved key interactors between both species and between cell types, with a subset of these proteins regulating RNA degradation and translation potentially relevant for the LINE-1 lifecycle itself. However, differences in experimental conditions in between studies could also influence this overlap. ORF1p interactors found in mouse spermatocytes ^35^ were also present in our analysis including CNOT10, CNOT11, PRKRA and FXR2 among others (Suppl_Table4). To unravel the physical interactions between the identified interactors of endogenous ORF1p within the mouse brain, we used the STRING database (Search Tool for Recurring Instances of Neighboring Genes, https://string-db.org/). This analysis generated a network representation, where physical interactions are represented by edges (Fig. 6D). In analogy with the GO term analysis, ORF1p displayed interactions with various clusters, including well-known RNA decapping complexes directed against LINE-1 RNA, which also encompassed DCP2 and DCP1A which had not previously been identified as interacting with ORF1p ^77^. Furthermore, ORF1p exhibited interactions with the SWI/SNF complex (highlighted in red) as well as subunits of the RNA polymerase II complex suggesting a direct or indirect association with accessible chromatin, a hitherto unknown interaction of ORF1p with chromatin compartments within the nucleus. Notably, a multitude of novel interactors belonged to the “neuronal cell body” and “neuron projections” clusters, proposing potential neuron-specific partners of ORF1p such as GRM2/5, BAI1, EPHA4, KCNN2, GRIK2 and DMD among others. A last cluster, formed by NCOA5 (Nuclear Receptor Coactivator 5), NXF1 (Nuclear RNA Export Factor 1), RANBP2 and NU133 (both nucleoporins), might imply a role for these interactions in L1-RNA nuclear export and/or a mechanism for the LINE-1 RNP to gain access to the nucleus in post-mitotic neurons. Altogether, the identification of known and novel interactors of ORF1p in the mouse brain suggests roles of ORF1p in the LINE-1 life cycle (RNA binding and metabolism, RNP formation, nuclear access) but also suggests potential novel physiological roles of ORF1p in the brain related to cytoskeleton organization, cGMP signaling, neuron-specific functions (i.e. synaptic signaling, Suppl_Table3) and chromatin organization and/or transcription regulation.

## Discussion

While LINE-1 derepression in aging has been extensively explored in peripheral tissues and various pathologies, including cancer, our understanding of LINE-1, particularly ORF1p, in the central nervous system remains limited ^20, 41, 78^. A recent search of ORF1p peptides in mass spectrometry data spanning 29 different healthy tissues did not reveal the presence of ORF1p in the brain, suggesting that its presence might lie below detection limits ^20^. Only a few studies explored and detected ORF1p expression in the brain, most focusing on a specific region (in mice ^17, 33^ in rats ^79^ and in human post-mortem brain ^19^), but it remained unclear if ORF1p is expressed throughout the entire brain, exhibits cell-type specificity, and most intriguingly, if its expression is influenced by the aging process. Here, using well characterized and validated antibodies, we demonstrate that ORF1p is expressed throughout the entire mouse brain and in at least three regions of the human post-mortem brain at steady-state. Leveraging a comprehensive workflow that incorporates brain atlas registration and machine learning algorithms, we quantified tens of thousands of brain cells, enabling a profound analysis of cell proportions, cell identities, densities and ORF1p expression levels across the entire brain. Surprisingly, more than one-fifth of detected cells expressed ORF1p. Regional variations in ORF1p expression levels were observed, with each region exhibiting distinct proportions, cell density, and signal intensity of ORF1p+ cells. In a non-neurologically diseased human brain, ORF1p was expressed in all three regions examined, that is the cingulate gyrus, the frontal cortex and the cerebellum. This is in accordance to an earlier study using histological staining, which found ORF1p expression in the human frontal cortex, the hippocampus, in basal ganglia, thalamus, midbrain and the spinal cord ^19^. This suggests, similarly to the mouse brain, a generalized expression across the human brain. On the transcriptomic level using long-read sequencing of GTEx tissues, brain and liver were highlighted as the organs displaying the highest expression of putatively active, full-length LINE-1 elements ^80^. However, when the authors looked at sub-regions, they found transcript expression in cerebellar hemispheres and the putamen, but not in the caudate and the anterior cingulate gyrus and frontal cortex ^80^. This is in contrast to our data and the data from Sur et al, where ORF1p was found to be expressed in the latter two regions using two different antibodies. We used the anti-human LINE-1 ORF1p antibody clone 4H1, a well characterized antibody ^78, 81^. While the sample size for the staining of human post-mortem tissues certainly needs to be increased in order to draw quantitative conclusions, the presence of the protein in two independent studies does point to a steady-state expression of ORF1p in the human brain.

It is interesting to note that ORF1p is expressed at steady-state in both, the mouse and human brain, despite the fact that evolutionary young and thus potentially ORF1-encoding LINE-1 elements in mice ^82^ (L1A, Tf/Gf) and in humans ^4^ (L1HS, L1PA2) differ significantly in number, sequence and regulation ^83^. Most differences lie in the 5’promoter region, but also ORF1 and ORF2 sequences are strikingly divergent between mouse and humans. For example, mouse LINE-1 promoters are composed of a varying number of monomers, a structure not found in human-specific LINE-1 promoters ^84^. This has obvious implications for LINE-1 expression regulation which can be very different, but examples of co-evolution of regulatory networks have been described ^85^ and might operate in the brain to regulate LINE-1 and thereby ORF1p expression.

In the mouse brain, we find ORF1p to be expressed predominantly if not exclusively in neurons using immunofluorescence and fluorescence-activated cell sorting (FACS) followed by Western blotting. This result is consistent with previous studies, such as the identification of ORF1p in excitatory neurons within the mouse frontal cortex ^86^, in parvalbumin neurons in the hippocampus ^18^, its presence in neurons in the ventral midbrain including in dopaminergic neurons ^17^ and the recognition of morphological similarities between stained neurons and ORF1p+ cells in a post-mortem hippocampus sample of a healthy individual ^19^. We also detected ORF1p in Purkinje cells in the mouse and human cerebellum. Neuronal specificity or preference of LINE-1 expression was also shown on the transcriptomic level in recent studies investigating LINE-1 expression in the mouse hippocampus, where neuronal LINE-1 expression exceeded that of astrocytes and microglia by approximately twofold ^20^, is abundant in parvalbumin interneurons ^18^ and single-nuclei RNA-seq data from the mouse hippocampus and frontal cortex which confirmed globally that repetitive elements including LINE-1 are more active in neurons than in glial cells ^86^. In the human brain, LINE-1 transcripts were found in greater quantities in neurons compared to non-neuronal cells by single-nucleus sequencing^87^. Furthermore, retrotransposition-competent LINE-1 elements were found expressed exclusively in neurons ^88^. While ORF1p expression is suggested to be expressed in microglia under experimental autoimmune encephalomyelitis conditions in the spinal cord ^89^, no evidence of such expression was observed in non-neuronal cells under non-pathological condition.

On average, throughout the mouse brain, the majority of neurons were positive for ORF1p and in some regions (i.e. the thalamus) around 80% of neurons expressed ORF1p. Comparing the results of both imaging approaches, the percentages of neurons expressing ORF1p in the ventral midbrain and frontal cortex were roughly similar (around 70% of neurons expressed ORF1p as quantified by confocal imaging and about 60% of neurons were identified as ORF1p+ using the slide scanner approach). In the human cingulate gyrus, we found that 37.2% of neurons express ORF1p and that 80% of cells expressing ORF1p were neurons, which are proportions similar to some regions of the mouse brain. It is however possible that these percentages are underestimated due to technical issues inherent to the machine-learning based algorithm for cell detection as our observations often indicated a positive signal in neurons which were classified as negative due to a particular shape or our stringent intensity threshold. A question which arises based on these findings is whether specific features distinguish ORF1p+ and ORF1p-neurons. One hint comes from a recent study suggesting that in the mouse hippocampus, it is the parvalbumin positive neurons that predominantly express ORF1p ^18^. However, as we show here, while PV-positive neurons often co-stain with ORF1p, not all ORF1p positive cells are PV-positive. In the mouse ventral midbrain, TH-positive dopaminergic neurons express higher levels of ORF1p compared to surrounding, non-dopaminergic neurons ^17^ (and this study Fig. 1B, panel 8). In the mouse cerebellum, ORF1p staining was detected in Purkinje cells and in the human post-mortem brain in Purkinje and possibly in Basket cells. Parvalbumin positive neurons are inhibitory neurons, so are Purkinje and Basket cells. However, dopaminergic neurons are modulatory neurons exerting excitatory and inhibitory effects depending on the brain region they act on. Specific neurons in the granular layer (i.e. Golgi and unipolar brush cells) of the cerebellum are inhibitory, but ORF1p negative, indicating that the decisive feature might not be the excitatory or inhibitory nature of a neuron. Further, ORF1p is expressed in excitatory (CamKIIa-positive) and CamKIIa-negative neurons in the mouse frontal cortex ^86^ and there is evidence of full-length L1 RNA expression in both excitatory and inhibitory neurons ^87^. While further studies are necessary to define the neuronal subtypes expressing ORF1p and their epigenetic make-up allowing this expression, it seems reasonable to conclude on the above-mentioned data that there is no neuronal sub-type specificity characterizing ORF1p expressing neurons. Another possibility is a cell-type specific chromatin organization permissive for the expression of LINE-1 and future single-cell studies in the mouse and human brain might reveal those differences.

Because transposable elements are known to become active in somatic tissues during aging ^15, 16, 28, 90, 91^, we aimed to investigate whether there was a corresponding increase at the protein level. In aged mice, ORF1p expression significantly increased throughout the mouse brain consistent with a previously documented increase in ORF1p outside the central nervous system in aged rats ^79, 92^ and aged mice ^90^ and in neurons of layer 2/3 of the mouse frontal cortex^86^. By quantifying the mean intensity of ORF1p in over 70 000 cells identified as ORF1p+, we were able to characterize the extent of this increase in each anatomical sub-region. Remarkably, apart from the isocortex which did not show any change, ORF1p expression increased in all other brain regions by 7% to 27%, indicating a generalized increase of ORF1p expression in neurons throughout the brain (13%). We did not detect any change in cell identity of ORF1p expressing cells, that is, ORF1p expression remained predominantly if not exclusively neuronal. Globally, there was also no change in ORF1p-positive, neuronal or non-neuronal cell densities in the aged mouse brain. Further investigations are necessary to investigate the underlying mechanism of the loss of ORF1p+ cells in the dorsal striatum in the aged mouse brain and to examine a possible relationship to the change of proportions of cells in the ventral midbrain, a structure which contains the *SNpc* which projects to the dorsal striatum and which is prone to LINE-1 driven neuronal degeneration ^17^. Thus, while ORF1p intensities per cell increase significantly in older mice in several brain regions, here is no global change in ORF1p+ cell numbers.

An increase of ORF1p might have several direct or indirect consequences on a cell or here, on a neuron. As ORF1p is translated from a polycistronic LINE-1 RNA together with ORF2p, albeit in much higher amounts (the estimated ratio ORF1p to ORF2p is 240:1 in non-native conditions) ^93^, it can be expected that a LINE-1 ribonucleotide particles are formed and ORF2-dependent cell toxicity in form of genomic instability ^17, 21^ and single-stranded cytoplasmic DNA triggered inflammation ^27, 28, 90^ might result. This has been shown in mouse dopaminergic neurons where oxidative stress induced LINE-1 causally contributed to neurodegeneration ^17^.

Neurodegeneration was partially prevented by anti-LINE-1 strategies among which NRTIs ^17^ and similar LINE-1 protein-dependent neuronal toxicity has been shown in drosophila^31, 32^ and the mouse cerebellum ^33^.

In order to test whether an increase in LINE-1 is a feature of human brain aging, we turned to a unique RNA-seq dataset of human laser-captured dopaminergic neurons of 41 individuals ranging from 38 to 99 years ^54^. In accordance with our focus on LINE-1 sequences which are full-length and coding, we developed a rationale to interrogate LINE-1 families with representatives that are coding (L1HS, L1PA2, multimappers; RepeatMasker) and to specifically investigate full-length LINE-1 elements that have intact open reading frames for ORF1p and ORF2p (unique reads; L1Basev2 ^4^). Indeed, we find an increase in L1HS and L1PA2 elements in individuals ≥65y as well as an increase in specific full-length LINE-1 elements but only a trend for increase of all full-length LINE-1 in sum in the elderly. This analysis has technical limitations inherent to transcriptomic analysis of repeat elements especially as it is based on short-read sequences and on a limited and disequilibrated number of individuals in both groups. Nevertheless, we tried to rule out several biases by demonstrating that mappability did not correlate with expression overall and used a combination of visualization, post-hoc power analysis and analysis of the mappability profile of each differentially expressed full-length LINE-1 locus. Interestingly, dysregulated full-length LINE-1 elements in aged dopaminergic neurons did not correspond to those identified in bladder cancer ^94^ indicating the intricate nature of this expression across tissues and pathological conditions. Overall, a slight net sum increase of UID transcripts/cell might be sufficient for the production of “above steady-state” levels ORF1p and ORF2p. Further, a dissociation of LINE-1 transcript and protein levels in aging has been observed recently in excitatory neurons of the mouse cortex. In the absence of transcriptional changes of LINE-1, protein levels of ORF1p were increased ^86^.

We can only speculate about the reason for an increase in ORF1p in the aged brain. A recent single-cell epigenome analysis of the mouse brain suggested a specific decay of heterochromatin in excitatory neurons of the mouse brain with age which was paralleled by an increase in ORF1p, albeit equally in excitatory and inhibitory neurons, again not indicating any dependency of ORF1p regulation on the excitatory or inhibitory nature of neurons ^86^. Chromatin and particularly heterochromatin disorganization are a primary hallmark of aging ^91^ but other repressive cellular pathways which control the LINE-1 life cycle might also fail with aging ^13^. Another possibility is a loss of accessibility of repressive factors to the LINE-1 promoter or an age-dependent decrease in their expression. Matrix correlation analysis of several known LINE-1 regulators, both positive and negative, revealed possible regulators of young LINE-1 sequences in human dopaminergic neurons. Despite known and most probable cell-type unspecific regulatory factors like the heterochromatin binding protein CBX5/HP1 ^58^ or the DNA repair proteins XRCC5 and XRCC6 ^56^, we identified the homeoprotein EN1 as negatively correlated with young LINE-1 elements including L1HS and L1PA2. EN1 is an essential protein for mouse dopaminergic neuronal survival ^57^ and binds, in its properties as a transcription factor, to the promoter of LINE-1 in mouse dopaminergic neurons ^17^. As EN1 is specifically expressed in dopaminergic neurons in the ventral midbrain, our findings suggest that EN1 controls LINE-1 expression in human dopaminergic neurons as well and serves as an example for a neuronal sub-type specific regulation of LINE-1. Although these proteins are known regulators of LINE-1, this correlative relationship awaits experimental validation. The heterogenous, brain-wide presence of ORF1p expression at steady-state is intriguing. In cancer cell lines or mouse spermatocytes, ORF1p interacts with several “host” proteins, some if not most of which are related to the LINE-1 life cycle. However, a profile of endogenous ORF1p interactors in the mouse brain might inform on possible other and organ-specific functions besides its binding to the LINE-1 RNA in “cis”^35^. Among the total 424 potential interactors of endogenous ORF1p in the mouse brain, 38 partners had been previously identified by mass spectrometry in human cancers, cancerous cell lines and mouse spermatocytes ^35, 67, 72–76^ (Suppl_Table4). This supports the validity of the list of ORF1p partners identified, although we cannot rule out the possibility that unspecific protein partners might be pulled down due to colocalization in the same subcellular compartment. GO term analysis contained expected categories like “P-body”, mRNA metabolism related categories and “ribonucleoprotein granule”. We also identified NXF1 as a protein partner of ORF1p, a protein found to interact with LINE-1 RNA related to its nuclear export ^95^. This suggests the conservation of key interactors probably essential for completing or repressing the LINE-1 life cycle in both species, despite the divergence of mouse and human ORF1p protein sequences ^96^. Along these lines, several ORF1p protein partners we identified might complete the list of post-transcriptional regulators implicated in LINE-1 silencing. Recent work conducted on human cancerous cell lines has demonstrated that MOV10 orchestrates the recruitment of DCP2 for LINE-1 RNA decapping ^77^. In our analysis, we identified DCP2 along with DCP1A, known to enhance the decapping activity of DCP2 ^97^, and DCP1b, a pivotal component of the mRNA decapping complex ^98^. Intriguingly, MOV10 was not detected in our mass spectrometry analysis, despite its established role in recruiting DCP2 and forming a complex with L1-RNP to mediate LINE-1 RNA decapping, as reported by Liu et al ^77^. However, we found two enhancers of mRNA decapping, EDC3 and EDC4, both core components of P-bodies, a membrane-less organelle known to contain L1-RNP ^65^. Multiple ubiquitin-ligase proteins were found although not appearing as a significantly enriched GO term. These results add to the picture of the post-transcriptional and translational control of ORF1p and suggest that these mechanisms, despite a steady-state expression, are operational in neurons. Further, several neuron-specific interactors were identified belonging to GO term categories “neuron projection” (75 proteins) and “neuronal cell body” (5 proteins), again pointing to the neuron-predominant expression of ORF1p in the mouse brain. Other interesting aspects were raised from this analysis. Among significantly enriched GO terms, several were related to the cytoskeleton, the functional consequences of which need to be determined in future studies. Our screen also identified PDE10A as an interactor of ORF1p in the mouse brain, a PDE almost exclusively expressed in medium spiny neurons of the striatum and a target for treatment of neurological diseases related to basal ganglia function like Huntington’s disease, schizophrenia and Tourette syndrome ^99^. Interestingly, PDE10A inhibition is related to beta-catenin signaling, another GO term which emerged from our screen ^100^. Finally, we found components of RNA polymerase II and the SWI/SNF complex as partners of ORF1p. This might further indicate that ORF1p has access to the nucleus in mouse brain neurons as described for other cells ^101, 102^, however a bias due to a post-lysis effect cannot be excluded. These findings give rise to intriguing questions regarding the potential function of ORF1p in neuron in health and disease as (i) ORF1p is widely distributed throughout the brain under normal physiological conditions, (ii) ORF1p shows a wide range expression levels within and in between regions, (iii) ORF1p is expressed predominantly if not exclusively in neurons, (iv) but not in all neurons and (v) interacts with proteins that might not directly relate to the LINE-1 life cycle, some of which are neuron-specific. In addition, physicochemical properties of ORF1p to form compacted nucleic acid - bound complexes with sequestration potential were shown ^96, 103^. Future loss-of-function studies should help to shed light on the necessity of ORF1p for neuronal functions if they exist. This data spurs the idea of a possible “physiological” function of ORF1p as an integrative protein with exapted function in neuronal homeostasis and a loss of restriction in the aged brain limiting LINE-1 expression to steady-state levels.

## Materials & Methods

### Animals

Swiss OF1 wild-type mice (Janvier) were housed on a 12h light/dark cycle with free access to water and food. Mice were sacrificed at 3-month or 16-month. Animal experiments were performed according to the EU directive 2010/63/EU.

### Mouse tissue dissection and protein extraction

Tissues were extracted from 3-month-old and 16-month-old Swiss/OF1 mice. Briefly, the two hemispheres were separated in ice cold PBS -/-. For each mouse, one hemisphere was rinsed and fixed in 4% PFA for 1h followed by 24h of incubation in 30% sucrose. Hemispheres were kept at -20°C until being sliced on a freezing microtome (Epredia, HM 450) with a 20µm thickness. The other hemisphere was dissected in ice cold PBS -/- 1X and 6 brain regions were rinsed, cut in small pieces and dissociated separately using a large (21G) to small gauge (27G) needle in RIPA lysis buffer for 5 min. Lysates were kept on ice for 25 min, were sonicated for 15 min and supernatants were collected after a 30 min centrifugation at 4°C at 14 000 rpm. Proteins were quantified and Laemmli buffer was added before boiling for 10min at 95°C to be used for Western Blot.

### Human Samples

Cerebellum, frontal cortex and cingulate gyrus human samples were provided by Neuro-CEB biobank from a 78-year-old brain-healthy male and conserved at -80°C.

### Human Samples pulverization and protein extraction

We used the dry pulverizer Cryoprep (Covaris) for pulverization of tissue blocs. Each sample was disposed in a liquid-nitrogen precooled Tissue-tube bag and dry cryo-pulverized with one impact at the maximum level. The pulverized brain sample was then weighed and resuspended in lysis buffer (mg/v) (0.32M sucrose, 5mM CaCl2, 3mM Mg(CH3COOH)2, 0.1mM EDTA, 10mM Tris-HCL pH8, 1mM DTT, 0.1% TritonX-100 and Protease Inhibitors), kept on ice for 30 min with gentle up-and-down pipetting until homogenization. We added 2X RIPA buffer (v/v) to totals fractions for 30 min on ice. We then sonicated samples 2 times for 15 min. AtlasSupernatants were collected after a 30 min centrifugation at 14 000 rpm at 4°C, proteins were quantified and Laemmli buffer was added to be used for Western Blot. All samples were boiled 10 min at 95°C to be used for Western Blot.

### Cell culture and siRNA delivery

LUHMES were purchased from ATCC (CRL2927). Cells were cultured on 50 μg/mL Poly-L-ornithine (Merck) and 1 μg/mL human plasma fibronectin (Sigma) coated flasks and cultured in Advanced DMEM/F12 (Gibco) added with 1% N-2 supplement (Gibco), GlutaMax (Gibco) and 40 ng/mL human recombinant FGF (Peprotech) at in 5 % CO2, 37C° incubator. Differentiation was initiated by adding to the media 1 µg/ml doxycycline (Sigma), 2 ng/ml recombinant human GDNF (Peprotech) and 1 mM cAMP (Sigma). Media was changed every two days. Experiences were performed on day 7 of differentiation. Cells were passaged less than 12 times

MN9D were purchased from Merck (SCC281). Cells were cultured on 1 mg/mL poly-lysine (Merck) coated flasks and cultured in DMEM High Glucose (Sigma, D5796) with 10% FBS (Gibco) in 5 % CO2, 37C° incubator. Over a period of 10 days, cells were differentiated by adding to the media 1 mM n-butyrate (Sigma) and 1 mM dibutyryl cAMP (Sigma). Media was changed every two days. Experiences were performed on day 10 of differentiation. Cells were passaged less than 15 times.

LUHMES were lipofected using RNAiMAX (Invitrogen) at day 3 of differentiation with 100 nM of siRNA. MN9D were lipofected using RNAiMAX (Invitrogen) at day 7 of differentiation with 100 nM of siRNA. The siRNA sequences used were as follows:

siRNA control: TAATGTATTGGAACGCATA

siRNA ORF1 (LUHMES): AAGAAGGCTTCAGACGATCAA

siRNA ORF1 (MN9D): CTATTACTCTGATACCTAAAC

LUHMES were scrapped at day 7 of differentiation and MN9D at 10 days of differentiation, in RIPA buffer (10mM Tris-HCl, pH 8.0; 150mM NaCl; 1mM EDTA; 1% Triton X-100; 0.1% Sodium Deoxycholate; 0.1% SDS). Laemmli buffer was added and samples were boiled 10 min at 95C° before being loaded on a gel.

### Western Blot

We used 1.5mm NuPAGE 4-12% Bis-Tris-Gel (Invitrogen™). Proteins samples (sorted mouse brain cells: 10000 cells/ µl -> 5 µl loaded; human brain lysates: 10µg; mouse brain lysates: 20µg) were loaded and gel migration was performed with NuPAGE™ MES SDS Running Buffer (Invitrogen™) for 45 minutes at 200mV. Gels were transferred onto a methanol activated PVDF membrane (Immobilon) in a buffer containing: Tris 25 mM, pH=8,3 and Glycine 192 mM, during 1h30 at 400 mA. Membranes were blocked 30 min with 5% milk in TBST (0,2% Tween 20, 150 mM NaCl, 10 mM Tris pH:8). The primary antibodies (mouse ORF1p antibody: abcam ab216324; human ORF1p antibody: abcam ab245249) were diluted in 5% milk in TBST, and membranes were incubated o/n at 4 C°. After 3 x 10 min washing in TBST, membranes were incubated for 1h30 with the respective secondary antibodies diluted at a concentration of 1/2000 in 5% milk TBST. Membranes were washed 3 x 10 min in TBST and were revealed by the LAS-4000 Fujifilm system using Clarity Western ECL Substrate (Bio Rad) or Maxi Clarity Western ECL Substrate (Bio Rad).

### Immunostaining

Sagittal mouse brains slices were fixed for 10min in PFA 4% and rinsed 3 times for 10min in PBS -/-. Slices were then incubated 20 min in glycine 100mM, washed 3 times for 5min in PBS and immersed in 10mM citrate pH 6 at 62°C during 45min for antigen retrieval. Slices were then immersed 3 times in PBS with Triton X-100 0.2% and incubated in blocking buffer for 1,5 h (PBS with Triton X-100 0.2% and FBS (10%) previously inactivated 20min at 56°C (Gibco, 16141061). Primary antibodies (ORF1p antibody: abcam ab216324; NeuN antibody: GeneTex GTX00837) were diluted (1/200 and 1/500 respectively) in blocking buffer and incubated with slices overnight at 4°C and then washed 3 times for 10 min with PBS. For validation, an in-house ORF1p antibody was used (09) (guinea pig, 1/200). For non-neuronal marker (GFAP antibody: Millipore AB5541; Iba1 antibody: GeneTex GTX101495; Sox9 antibody: RnDsystems AF3075; S100β antibody: Sigma S2532), antibodies were diluted at 1/500. Additionally, WFA (Sigma L1516) and PV antibodies (Swant PV235) were used, diluted at 1/500 and 1/1000 respectively. Suitable secondary antibodies (Invitrogen) and Hoechst (Invitrogen, 15586276) were incubated for 1,5h at 1/2000 in PBS with inactivated FBS (10%) and washed 3 times 10 min in PBS. To quench tissue autofluorescence, especially lipofuscin, TrueBlack Plus (Biotium) in PBS was used during 10min. Slices were rinsed 3 times in PBS and mounted with Fluoromount (Invitrogen™).

For human cingulate gyrus stainings, the same protocol was performed, with the difference that a human ORF1p antibody (Abcam 245249) was used. Mouse and human brain slices were imaged by the Axioscan 7 Digital Slide Scanner (Zeiss) or a Spinning Disk W1 confocal microscope (Yogogawa).

### Blocking peptide

The ORF1p antibody (abcam ab216324) was incubated 2h on a turning wheel with excess (4:1) of mouse ORF1p recombinant protein as in ^17^ before the blocked antibody was used in the above-described immunofluorescence protocol.

### Quantification of confocal acquisitions

Analysis was conducted using a custom-written plugin developed for the Fiji software, incorporating Bio-Formats ^104^ and 3D ImageJ Suite ^105^ libraries. Code is freely available online at https://github.com/orion-cirb/DAPI_NEUN_ORF1P. Nuclei were detected in the Hoechst channel downscaled by a factor of 2 with the 2D-stitched version of Cellpose ^106^ (percentile normalization = [1-99], model = ‘cyto’, diameter = 30, flow threshold = 0.4, cell probability threshold = 0.0, stitching threshold = 0.75). The segmented image was then rescaled to its original size, and the obtained 3D nuclei were filtered by volume to reduce false positive detections. NeuN+ and ORF1p+ cells were detected in their respective channel using the same approach as for nuclei detection, but with adjusted Cellpose settings (model = ‘cyto2’, diameter = 40, flow threshold = 0.4, cell probability threshold = 0.0, stitching threshold = 0.75). Finally, each cell was associated with a nucleus having at least half of its volume in contact with. Cells without any associated nucleus were discarded. Each nucleus was thus labeled according to NeuN and/or ORF1p positivity.

Nuclei and cell detection using the respective Cellpose models and hyperparameters were evaluated on eight images per channel, capturing intensity variability across different mouse ages and brain regions. A total of approximately 2,000 nuclei and 1,000 NeuN and ORF1p cells were manually annotated. We evaluated model performance with the average precision (AP) metric, computed from the number of true positives (TP), false positives (FP), and false negatives (FN) as 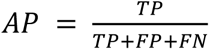 at the commonly used Intersection over Union (IoU) threshold of 0.5. The AP at an IoU threshold of 0.5 was 0.995 for nuclei, 0.960 for NeuN, and 0.974 for ORF1p cells. These results confirmed that the selected models and hyperparameters were well-suited for processing the entire dataset.

### ABBA Registration and Qupath analysis

Each sagittal brain section was registered with the Allen Mouse Brain Atlas (CCFv3 ^107^) using the Aligning Big Brains & Atlases plugin ^34, 108^ in Fiji. Slices were first manually positioned and oriented along the Z-axis. Automated affine registration was then applied in the XY plane, followed by manual refinement. The final registration results were imported into QuPath software ^109^ for downstream processing. Cell analysis in each brain subregion was performed with custom Groovy scripts developed for QuPath. Code is freely available online at https://github.com/orion-cirb/QuPath_ORF1P. Hoechst nuclei were detected with StarDist 2D ^110^, applying the DSB 2018 pretrained model with the following parameters: percentile normalization = [1-99], probability threshold = 0.82, overlap threshold = 0.25. Cells in NeuN and ORF1p channels were detected with Cellpose 2D (percentile normalization = [1-99], model = ‘cyto2’, diameter = 30, flow threshold = 0.4, cell probability threshold = 0.0). Nuclei and cells were then filtered by area and intensity to minimize false positive detections. Minimum intensity threshold was based on the channel background noise, which was estimated for each subregion as the mean intensity of pixels not belonging to any detected nucleus or cell in the respective channel. Finally, each cell was associated with a nucleus having its centroid located inside the cell mask. Cells without an assigned nucleus were discarded, cells with associated nuclei were classified as NeuN+ or NeuN- and ORF1p+ or ORF1p-. Intensity values were normalized by subtracting the background noise computed in the corresponding channel and subregion. As a last step, subregional results were merged into regional ones and data were analyzed using the Pandas Python library ^111^.

Nuclei and cell detection were evaluated on 14 images per channel, corresponding to approximately 800 nuclei and 400 NeuN and ORF1p cells manually annotated. The average precision (AP) at an IoU threshold of 0.5 was lower than for confocal images: 0.806 for nuclei, 0.675 for NeuN, and 0.695 for ORF1p cells. This decline in performance was primarily due to a lower signal-to-noise ratio in slide scanner images, leading to an increased number of false positives and false negatives. While fine-tuning the models could enhance detection robustness, the selected models and hyperparameters were considered suitable for processing the entire dataset.

### FACS

Mouse brains were dissociated with Adult Brain Dissociation kit (Miltenyi Biotec, 130-107-677) and incubated with the coupled antibody NeuN Alexa 647 (Abcam, ab190565) or the control isotype IgG Alexa 647 (Abcam, ab199093). Stained cells were filtered a last time with a 40µm filter before FACS sorting (FACS ARIA II). Neuronal and non-neuronal cells were separately collected in PBS -/- 2m EDTA and then centrifugate (5min at 700rpm). Pellets were resuspended in RIPA for protein extraction in an appropriate volume in order to achieve equal cell concentrations (10 000 cells/µl).

### RNA-seq analysis

The RNA-seq dataset from Dong et al ^54^ was downloaded from dbGAP (phs001556.v1.p1) and contains unstranded paired-end 50bp and 75bp reads from pooled laser-capture micro-dissected dopaminergic neurons from human post-mortem brain (107 samples) from 93 individuals w/o brain disease. RNA-seq had been done on total and linearly amplified RNA. We focused our analysis on data obtained with 50bp reads, in order to avoid mappability bias, while still regrouping all age categories (n=41; with ages ranging from 38 to 97 (mean age: 79.88 (SD ±12.07); n=6 ≤65y; n=35 >65y; mean PMI: 7.07 (SD ±7.84), mean RIN: 7.09 (±0.94)). Sequencing reads were aligned on the Human reference genome (hg38) using the STAR mapper (v2.7.0a) 3 and two different sets of parameters. Genome-wide individual repeat quantification was performed using uniquely mapped reads and the following STAR parameters: --outFilter mapNmax 1–-alignEndsType EndToEnd–-outFilterMismatchNmax 999–-outFilterMismatchNoverLmax 0.06. Repeats class, family and name quantification was performed using a random mapping procedure and the following parameters : -- outFilterMultimapNmax 5000–-outSAMmultNmax 1–-alignEndsType EndToEnd–-outFilterMismatchNmax 999–-outFilterMismatchNoverLmax 0.06–-outSAMprimaryFlag OneBestScore–-outMultimapperOrder Random. Repeats annotations were downloaded from the UCSC Table Browser (repeatMasker database: https://genome.ucsc.edu/cgi-bin/hgTables) and coordinates of LINE-1 full length and coding elements were downloaded from the L1base database 2 (http://l1base.charite.de/l1base.php; ^4^) selecting LINE-1 full length elements containing two predicted complete open reading frames for ORF1 and ORF2 (UID= Unique IDentifier) from the LINE-1 database (http://l1base.charite.de/l1base.php) and corrected genomic intervals with the repeat masker annotation of the corresponding genomic locus. Repeat quantification from the aligned data was done using a gtf file composed of all genes (Gencode v29) and all individual repeat elements. This strategy was used to avoid overestimation of repeat elements due to overlaps with expressed genes. For individual repeat quantification of the full length L1 elements (L1base), we therefore used a gtf of all genes and all L1base entries, and ran the FeatureCounts tool ^112^ with the following parameters: -g gene_id -s 0 -p. In the context of the family-based analysis, we used a gtf with all genes and all annotated repeats elements and ran FeatureCounts with -g gene_family -s 0 -p -M. Before DeSeq2 analysis, we remove all genes and repeat elements with less than 10 reads in a minimum of n individuals, n being the number of individuals in the condition containing the fewest individuals (“young” condition: n=6, 38-65y, mean 57.5 years). We use the same conditions with genes and UIDs with less than 3 reads in a minimum of n individuals. Finally, we calculated the scaling factors using DeSeq2 on all genes + all repeat elements or all gene + UID according to the quantification method and then applied these scaling factors to the corresponding counts tables.

In order to test for the mappability of each UID (= full-length and coding LINE-1), we extracted the bed track « main on human:umap50 (genome hg38) from the UCSC genome browser (≈ 7Mio regions) directly into Galaxy (usegalaxy.org) and joined genomic intervals with a minimum overlap of 45bp of this dataset with a dataset containing the annotation of UIDs extracted from L1Basev2 ^4^ corrected in length with repeat masker and completed with information on whether the UID is intra- or intergenic and, if intragenic, in which gene (NM_ID, chr, strand, start, end, gene length, number of exons, gene symbol) the fl-LINE-1 is located, which resulted in 1266 regions. We then used the “group on data and group by” function in Galaxy and counted the number of overlapping 50kmers with all 146 UIDs (=mappability score). Correlation analysis (non-parametric Spearman) was then done between the mappability score and the normalized read counts.

For visualization of expression, bigwig files were generated for each age group, that is </=65y and >65y, respectively. We used bamcoverage to obtain bigwig files (normalized by cpm). Then for each age group an average bigwig was generated using bigwigAverage from deeptools (galaxy version 3.5.4); Bigwigs were loaded into IGV alongside tracks showing mappability (Umap50) and selected tracks from repeat masker (LINE, SINE and LTR).

Post-hoc power analysis was performed using the “Post-hoc Power Calculator” (https://clincalc.com/stats/Power.aspx).

### Immpunoprecipitation of ORF1p from the mouse brain

For immunoprecipitation, we used ORF1p (abcam, ab245122) and IgG rabbit (abcam, ab172730) antibodies. The antibodies were coupled to magnetic beads using the Dynabeads® Antibody Coupling Kit (Invitogen, 14311D) according to the manufacture’s recommendations. We used 5µg of antibody for 1 mg of beads and used 1.5mg of beads for IP. Individual mouse brain lysates (n=5), homogenized using dounce and sonicated, were incubated with ORF1p or IgG-control coupled beads and a small fraction was kept as input. Each of these two tubes containing coupled beads and brain lysates were diluted in 5 ml buffer (10 mM Tris HCl, 150 mM NaCl, protease inhibitor). The samples were then incubated overnight on a wheel at 4°C. Samples were then washed 3 times with 1 ml buffer (10 mM Tris HCl pH 8, 200 mM NaCl) using a magnet and then resuspended in the same buffer. The samples were boiled in Laemmli buffer (95°C, 10 min) and 20 µl of each sample were loaded on a 4-12% Nupage gel (Invitrogen, NP0336) to be revealed by WB. For samples used in Mass Spectrometry study, beads were washed with buffer (10 mM Tris HCl pH 8, 200 mM NaCl) using a magnet. After 3 washes with 1ml buffer the beads were washed twice with 100 µL of 25 mM NH4HCO3 (ABC buffer). Finally, beads were resuspended in 100 μl of 25mM ABC buffer and digested by adding 0.20 μg of trypsine/LysC (Promega) for 1 hour at 37 °C. A second round of digestion was applied simultaneously on the beads by adding 100 µL of 25 mM ABC buffer and to the previous digest by adding 0.20 µg of trypsin/LysC for 1 hour at 37 °C. Samples were then loaded into homemade C18 StageTips packed by stacking three AttractSPE® disk (#SPE-Disks-Bio-C18-100.47.20 Affinisep) into a 200 µL micropipette tip for desalting. Peptides were eluted using a ratio of 40:60 CH3CN:H2O + 0.1% formic acid and vacuum concentrated to dryness with a SpeedVac device. Peptides were reconstituted in 10 µL of injection buffer in 0.3% trifluoroacetic acid (TFA) before liquid chromatography-tandem mass spectrometry (LC-MS/MS) analysis.

### Immunoprecipitation of ORF1p from LUHMES cells (human) differentiated into mature dopaminergic neurons

For immunoprecipitation, we used ORF1p (Millipore, MABC 1152) and IgG mouse (Thermofisher, #31903) antibodies. The antibodies were coupled to magnetic beads using the Dynabeads® Antibody Coupling Kit (Invitogen, 14311D) according to the manufacturer’s recommendations. We used 8 µg of antibody for 1 mg of beads. The appropriate volume of buffer was added to the coupled beads to achieve a final concentration of 10 mg/ml. Cells were washed with 1X PBS and harvested using 1 ml of lysis buffer (10 mM Tris HCl pH 8, 150 mM NaCl, NP40 0.5% v/v, protease inhibitor 10 µl/ml). Samples were sonicated for 15 min at 4°C, then centrifuged at 1200 rpm for 15 min at 4°C. The supernatants obtained were transferred to a new microcentrifuge tube and then separated into 3 tubes. One tube was used as the input and the two other two tubes were used for the control and ORF1p IP, respectively. Each of these two tubes were then diluted to 1.5 ml with buffer (10 mM Tris HCl pH8, 150 mM NaCl, 10 µl/ml protease inhibitor) to dilute the NP40. The samples were then incubated overnight on a wheel at 4°C, then washed 3 times (first wash corresponds to post-bead samples) with 1 ml buffer (10 mM Tris HCl pH 8, 150 mM NaCl, 10 µl/ml protease inhibitor) using a magnet and then resuspended in the same buffer. The samples were boiled in Laemmli buffer (95°C, 10 min) and 20 µl of each sample were deposited on a 4-12% Nupage gel (Invitrogen, NP0336).

### Mass Spectrometry

Online chromatography was performed with an RSLCnano system (Ultimate 3000, Thermo Scientific) coupled to a Q Exactive HF-X with a Nanospay Flex ion source (Thermo Scientific). Peptides were first trapped on a C18 column (75 μm inner diameter × 2 cm; nanoViper Acclaim PepMap^TM^ 100, Thermo Scientific) with buffer A (2/98 MeCN/H2O in 0.1% formic acid) at a flow rate of 2.5 µL/min over 4 min. Separation was then performed on a 50 cm x 75 μm C18 column (nanoViper Acclaim PepMap^TM^ RSLC, 2 μm, 100Å, Thermo Scientific) regulated to a temperature of 50°C with a linear gradient of 2% to 30% buffer B (100% MeCN in 0.1% formic acid) at a flow rate of 300 nL/min over 91 min. MS full scans were performed in the ultrahigh-field Orbitrap mass analyzer in ranges m/z 375–1500 with a resolution of 120 000 at m/z 200. The top 20 intense ions were subjected to Orbitrap for further fragmentation via high energy collision dissociation (HCD) activation and a resolution of 15 000 with the intensity threshold kept at 1.3 x 105. We selected ions with charge state from 2+ to 6+ for screening. Normalized collision energy (NCE) was set at 27 and the dynamic exclusion of 40s. For identification, the data were searched against the Mus Musculus (UP000000589_10090 012019) Uniprot database using Sequest HT through proteome discoverer (version 2.4). Enzyme specificity was set to trypsin and a maximum of two-missed cleavage sites were allowed. Oxidized methionine, Met-loss, Met-loss-Acetyl and N-terminal acetylation were set as variable modifications. Maximum allowed mass deviation was set to 10 ppm for monoisotopic precursor ions and 0.02 Da for

MS/MS peaks. The resulting files were further processed using myProMS ^113^ v3.10.0. FDR calculation used Percolator and was set to 1% at the peptide level for the whole study. The label free quantification was performed by peptide Extracted Ion Chromatograms (XICs), reextracted by conditions and computed with MassChroQ version 2.2.21 ^114^. For protein quantification, XICs from proteotypic peptides shared between compared conditions (TopN matching) with missed cleavages were used. Median and scale normalization at peptide level was applied on the total signal to correct the XICs for each biological replicate (n=5). To estimate the significance of the change in protein abundance, a linear model (adjusted on peptides and biological replicates) was performed, and p-values were adjusted using the Benjamini– Hochberg FDR procedure. Proteins with at least three peptides, identified in each biological replicates of ORF1p condition, a 10-fold enrichment and an adjusted p-value ≤ 0.05 were considered significantly enriched in sample comparisons. Unique proteins were considered with at least three peptides in all replicates. Protein selected with these criteria were used for Gene Ontology enrichment analysis and string network analysis.

The mass spectrometry proteomics data have been deposited to the ProteomeXchange Consorti-um (http://proteomecentral.proteomexchange.org) via the PRIDE partner repository ^115^ with the dataset identifier PXD047160.

### GO term and STRING network analysis

Gene Ontology analysis was performed using GO PANTHER ^116^ and String network physical interactions were retrieved using the STRING database v11.5 (https://string-db.org/) and then implemented in Cytoscape software ^117^.

### Statistical analyses

In column comparisons, data in each column were tested for normality using two normality and lognormality tests (Shapiro-Wilk test and Kolmogorov-Smirnov test). Data which passed the normality tests were analyzed subsequently by a parametric test, data which did not pass the normality tests were analyzed by a non-parametric statistical test as indicated in the figure legends. The significance threshold was defined as p<0.05 except stated otherwise. Statistical analyses were done with PRISM software (v10).

## Acknowledgments

This work was supported by the Fondation de France (00086320, to J.F.), the Fondation du Collège de France (to J.F. and T.B.), the Fondation NRJ/Institut de France (to J.F.), the Fondation Alzheimer and the National French Agency for Research (ANR-20-CE16-0022 NEURAGE). We gratefully acknowledge the Orion Technological Core (IMACHEM-IBiSA) of CIRB for their support, member of the France-BioImaging research infrastructure, especially Estelle Anceaume and Julien Dumont for assistance with slide scanner and spinning disk acquisition and Magali Fradet for assistance with FACS analysis. We also thank the Fondation Bettencourt Schueller for their support.

## Contributions

T.B. carried out most of the experimental work and analyzed the data. T.B., S.S. and J.F. analyzed the transcriptomic data. S.S participated in the experimental work and established and performed the loss of function experiments. O.M.B. performed the FACS analysis. T.B. and J.F. wrote the manuscript. T.B., P.M. and H.M. developed the image analysis pipeline. B.L. carried out the MS experimental work and D.L. supervised MS and performed data analysis with T.B.. Read alignment, quality control and mapping were done by N.S.. J.F. and R.L.J. conceived and supervised the project.

## Supplementary Figures

**Suppl Fig. 1.**
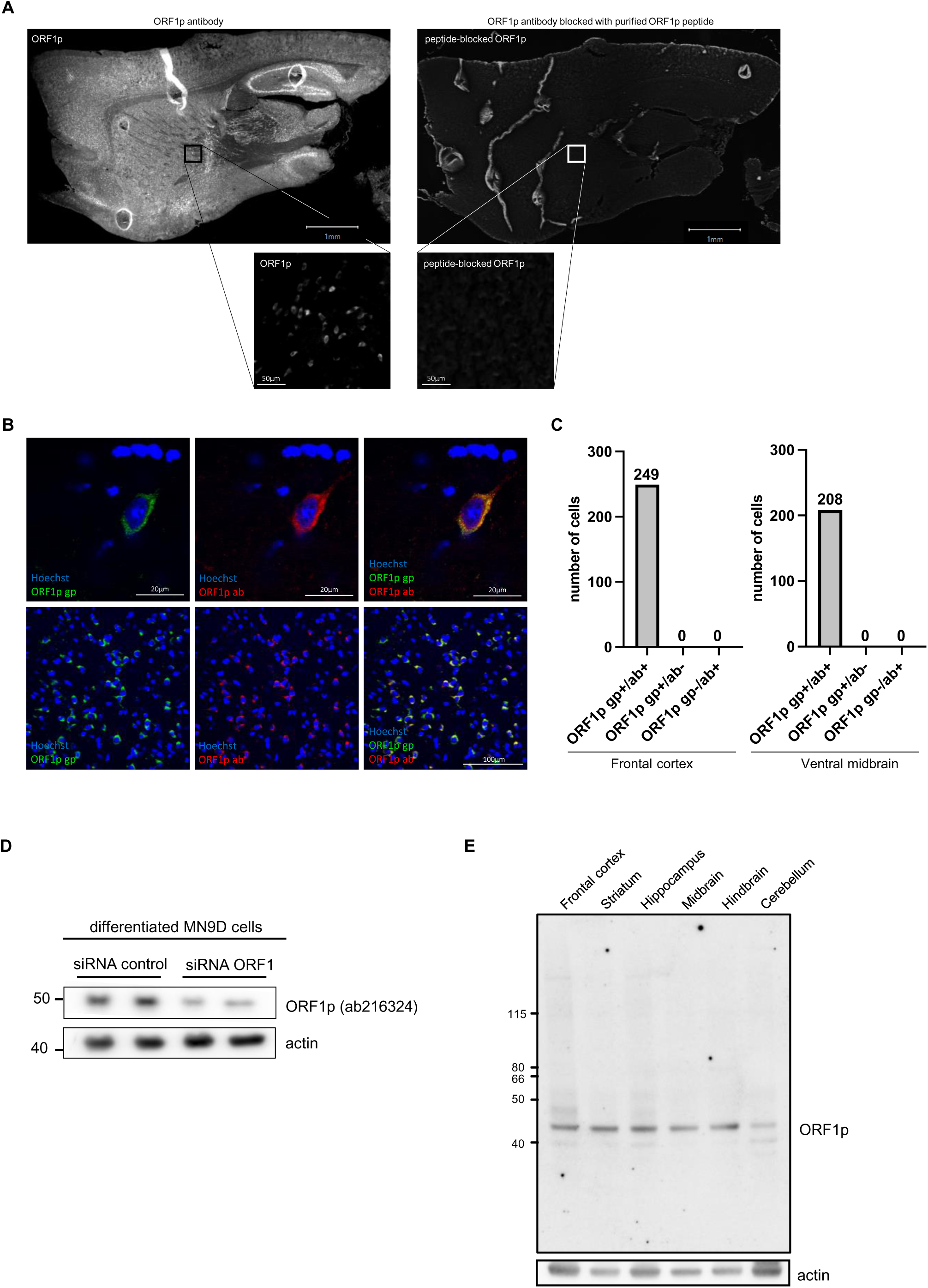
(A) Specific antigen recognition by the ORF1p antibody. IHC showing ORF1p positives cells in sagittal mouse brain slice (left) and abolition of the signal when blocking the antibody with purified ORF1p (right). (B) Representative acquisition showing ORF1p stainings obtained with a commercially available ORF1p antibody (abcam ab216324) used in this study (red) and with an in-house ORF1p gp antibody (guinea pig, green) in the mouse brain. Scalebar = 20µm (top) and 100µm (bottom). (C) Quantification of double positive cells (gp+/ab+) using the commercial ORF1p ab antibody (abcam ab216324) and in-house ORF1p gp antibody (guinea pig) versus single-positive cells (gp+/ab- and gp-/ab+) in mouse frontal cortex (left) and ventral midbrain (right). (D) Validation of the ORF1p antibody (abcam ab216324) by knock-down of ORF1p in mouse dopaminergic neurons in cluture. MN9D cells differentiated into mature mouse dopaminergic neurons were transfected with either control siRNA or ORF1p siRNA and ORF1p was revealed by Western blot. Actin was used as a loading control. (E) ORF1p is expressed in six different brain regions in the mouse. Brain regions were micro-dissected from a three-month old mouse brain. Western blot showing ORF1p expression in 6 brain regions. ORF1p (Top), Actin (bottom).

**Suppl Fig. 2.**
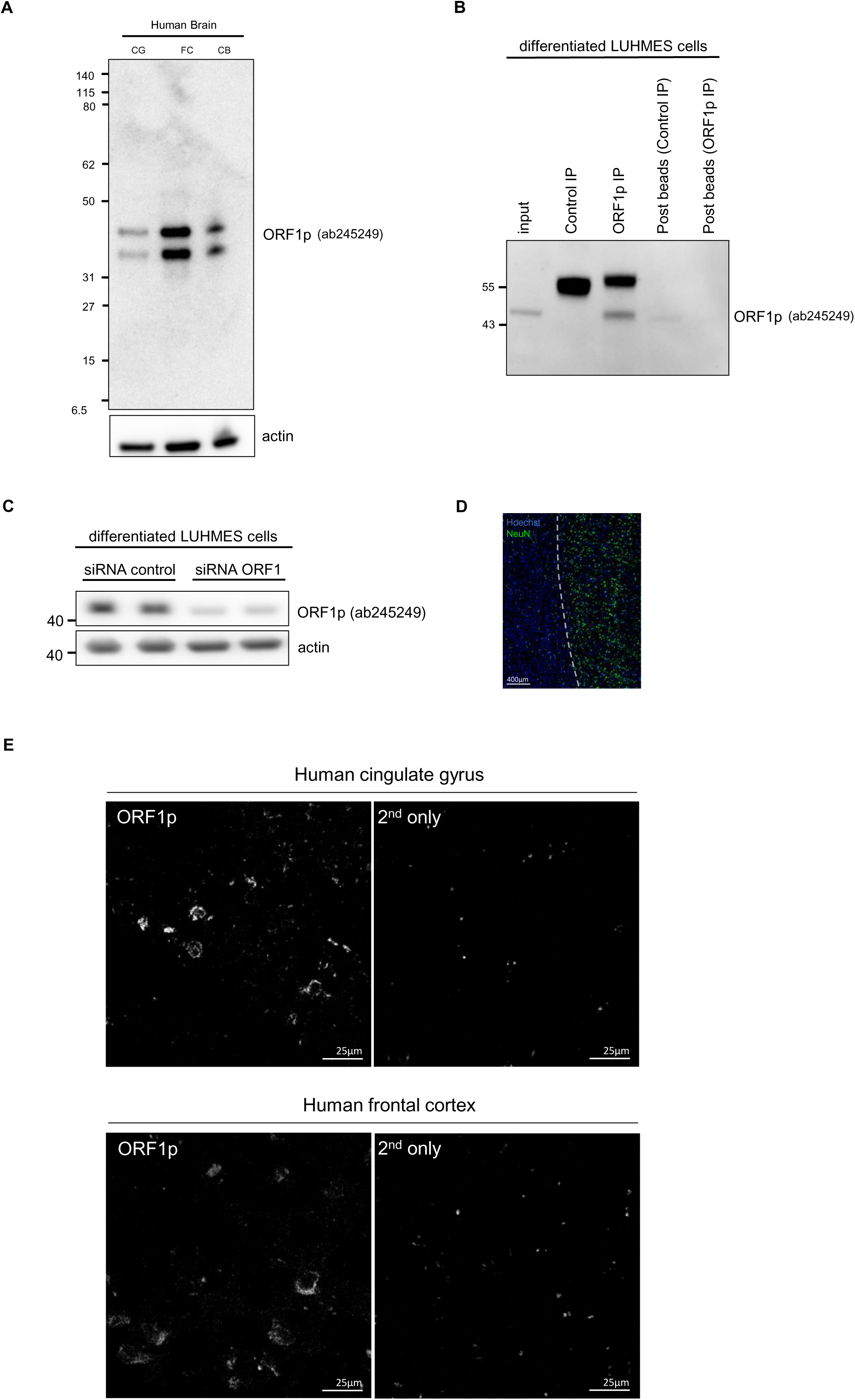
(A) ORF1p is expressed in the human post-mortem brain. Complete membrane of the ORF1p Western blot on human post-mortem cingulate gyrus (CG), frontal cortex (FC) and cerebellum (CB) shown in Fig. 1J. (B) Immunoprecipitation of ORF1p from LUHMES cells which were differentiated into mature human dopaminergic neurons. First lane: Input; second lane: Control IP; third lane: ORF1p IP; forth lane: post-beads from control IP; fifth lane: post-beads from ORF1p IP. (C) Validation of the human ORF1p antibody by knock-down. LUHMES cells differentiated into mature human dopaminergic neurons were transfected with either control siRNA or ORF1p siRNA and ORF1p was revealed by Western blot using the ORF1p abcam ab245249. Actin was used as a loading control. (D) Representative slide-scanner acquisition of a human cingulate gyrus section showing NeuN positives cells (green) mostly located in the grey matter (right) compared to the white matter from a brain-healthy individual; scale bar = 400µm. A zoom into the grey matter region showing ORF1p stainings is shown in Figure 2H. (E) Control experiment using a 2^nd^ antibody-only control on human cingulate gyrus and frontal cortex alongside ORF1p stainings. Representative confocal microscopy images showing ORF1p or only secondary antibody. z-projection; scalebar = 25µm.

**Suppl Fig. 3.**
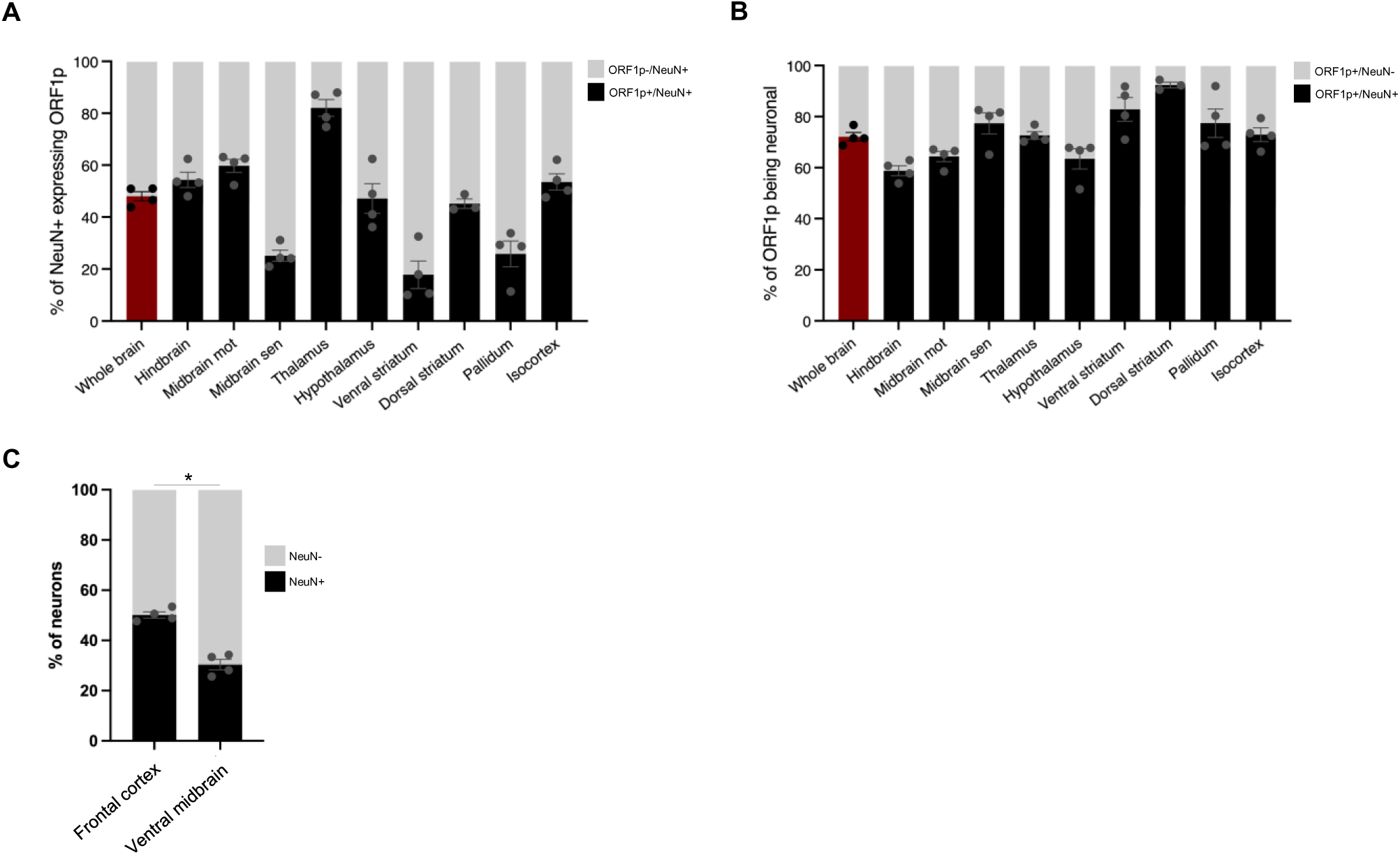
(A) Proportion of neurons (NeuN+) expressing ORF1p in the whole brain (left, red) and in different regions of the mouse brain (black bars) or not (grey) as quantified using the cell detection pipeline on large-scale images. (B) ORF1p cell identity. Proportion of ORF1p+ cells identified as NeuN+ (black) or NeuN- (grey), in the whole brain (left, red)) and in 9 different regions analyzed (right) using the cell detection pipeline on large-scale images presented in Figure 1A; data is represented as mean ± SEM, n=4 mice. (C) Proportion of neurons in the frontal cortex and ventral midbrain quantified using the confocal approach. *p<0.05, chi-square test on the cell number of the different cell-types analyzed; n=4 mice, data is represented as mean ± SEM.

**Suppl Fig. 4.**
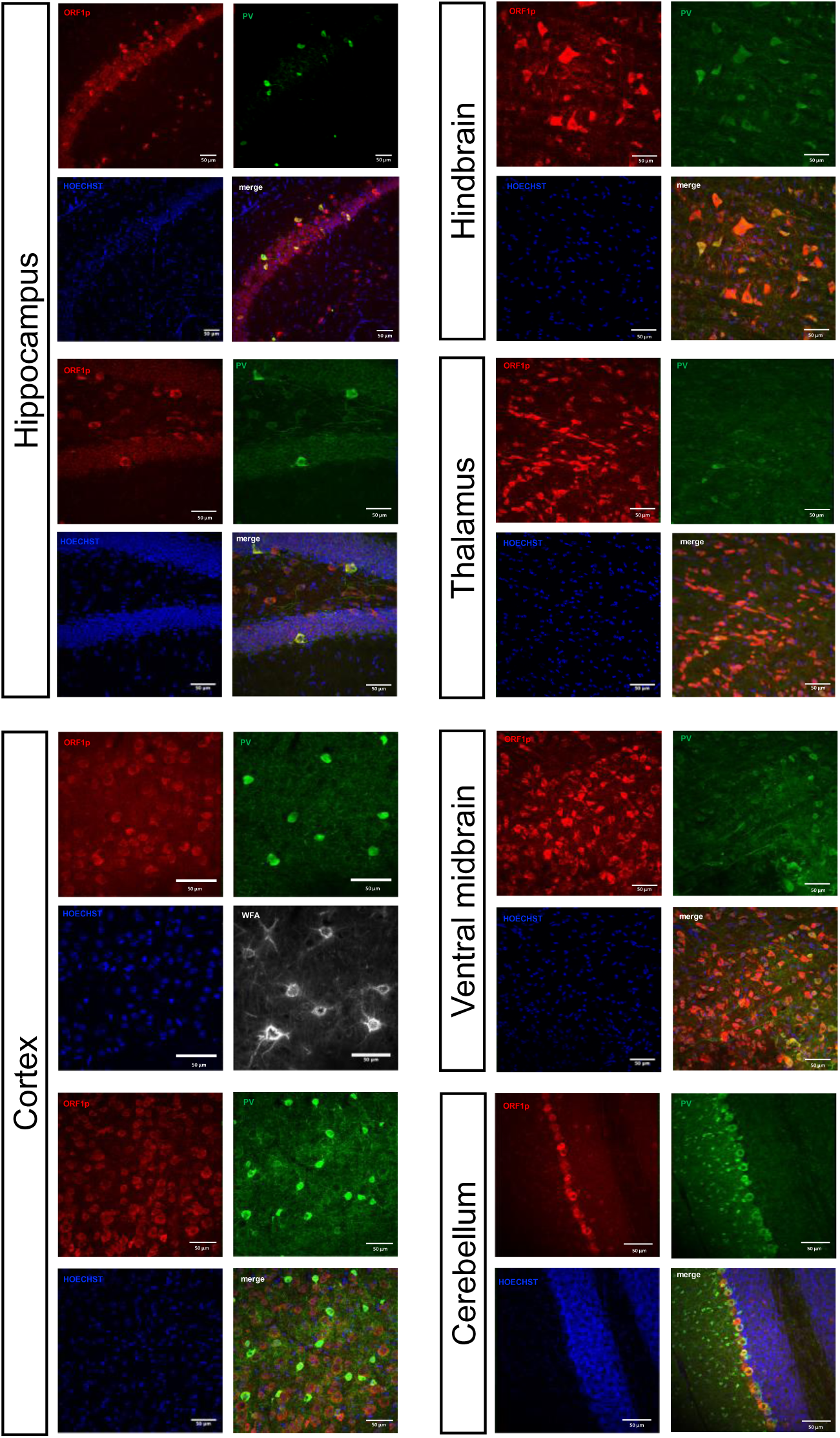
Representative confocal microscopy images showing ORF1p (red), PV (green), WFA (grey), Hoechst (blue) and merged images in six regions of the mouse brain, scalebar = 50µm.

**Suppl Fig. 5.**
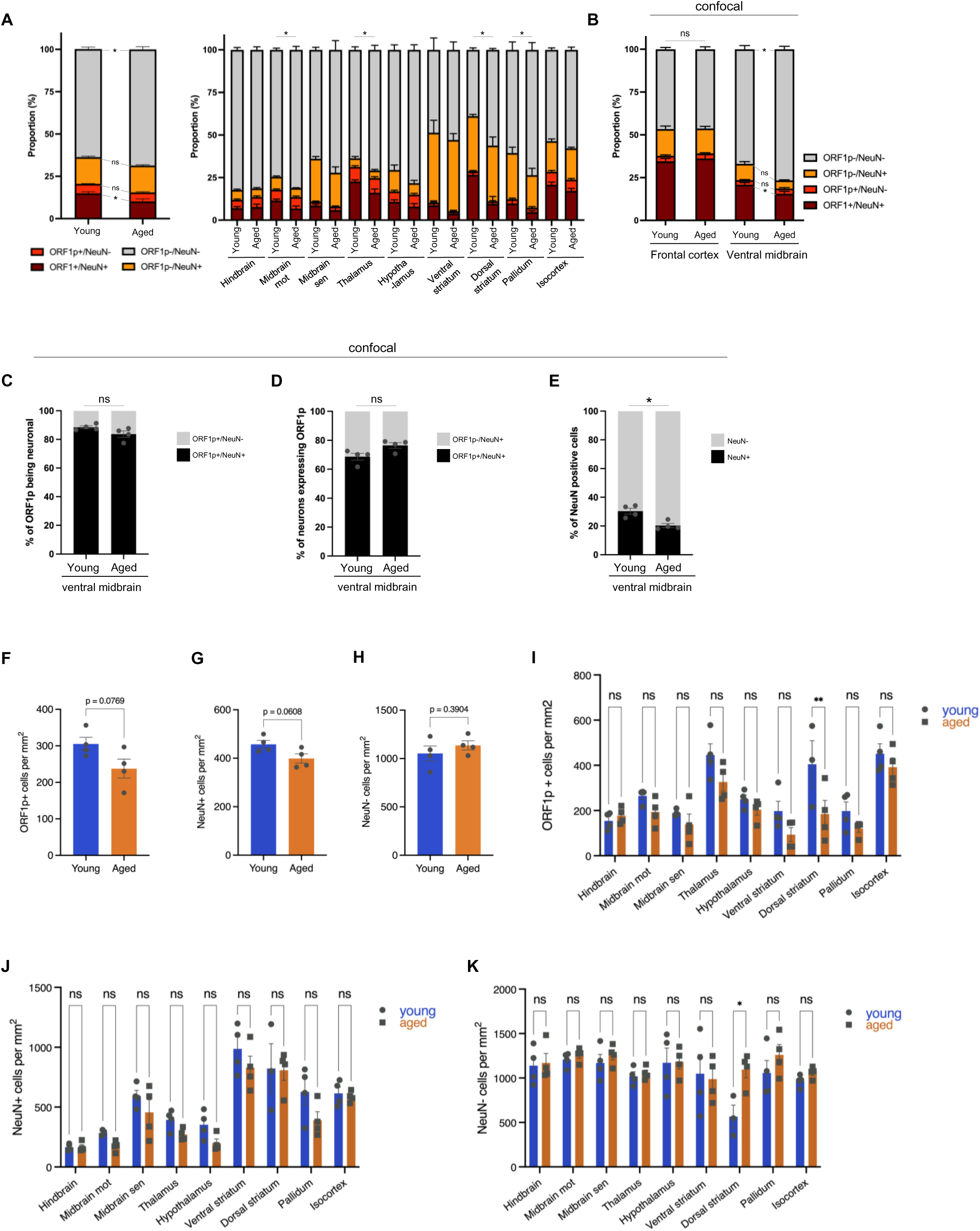
(A) Proportion of ORF1p+/NeuN+, ORF1p+/NeuN+, ORF1p+/NeuN-, ORF1p-/NeuN+ and ORF1p-/NeuN-cell-type in the whole brain (left) and the different analyzed regions (right) of young (three-month aged, n=4) and aged (16-month-old, n=4) mice using the cell detection pipeline on large scale images presented in Figure 1A, data is represented as mean ± SEM. Exact values can be found in Suppl_Table1. (B) Proportion of ORF1p+/NeuN+, ORF1p+/NeuN+, ORF1p+/NeuN-, ORF1p-/NeuN+ and ORF1p-/NeuN-cell-type in two different regions of young and aged mice, analyzed on multiple z-stack confocal images. ns: non-significant; *p<0.05 calculated using two-way ANOVA with sidak’s multiple comparisons test on the cell number of the different cell-types analyzed; data is represented as mean ± SEM. (C) Proportion of ORF1p+ cells being neuronal in the ventral midbrain comparing young and aged mice as quantified using the confocal approach. Kolmogorov-Smirnov test; data is represented as mean ± SEM. (D) Proportion of neurons expressing ORF1p in the ventral midbrain comparing young and aged mice as quantified using the confocal approach. Kolmogorov-Smirnov test; data is represented as mean ± SEM. (E) Proportion of neurons in the ventral midbrain comparing young and aged mice as quantified using the confocal approach. Kolmogorov-Smirnov test; data is represented as mean ± SEM. (F) ORF1p+ cell density in the brain comparing young and aged mice, analyzed on large-scale images. ORF1p+ cell number per mm2 in young and aged mice (n=4 per condition); Shapiro Wilk normality test followed by an unpaired, two-tailed t-test. (G) Neuronal cell density in the brain comparing young and aged mice, analyzed on large-scale images. NeuN+ cells per mm2 in young and aged mice (n=4 per condition); Shapiro Wilk normality test followed by an unpaired, two-tailed t-test. (H) Non-neuronal cell density in the brain comparing young and aged mice, analyzed on large-scale images. NeuN- cells per mm2 in young and aged mice (n=4 per condition); Shapiro Wilk normality test followed by an unpaired, two-tailed t-test. (I) ORF1p+ cell density in young and aged mice in nine brain regions, analyzed on large-scale images. Data is represented as mean ± SEM, n=4 mice per condition. Two-way ANOVA followed by a Sidak’s multiple comparisons test, alpha = 0.05. Adjusted p-values <0.05 are shown. (J) NeuN+ cell density in young and aged mice in nine brain regions, analyzed on large-scale images. Data is represented as mean ± SEM, n=4 mice per condition. Two-way ANOVA followed by a Sidak’s multiple comparisons test, alpha = 0.05. Adjusted p-values <0.05 are shown. (K) Non-neuronal cell density in young and aged mice in nine brain regions, analyzed on large-scale images. Data is represented as mean ± SEM, n=4 mice per condition. Two-way ANOVA followed by a Sidak’s multiple comparisons test, alpha = 0.05. Adjusted p-values <0.05 are shown.

**Suppl Fig. 6.**
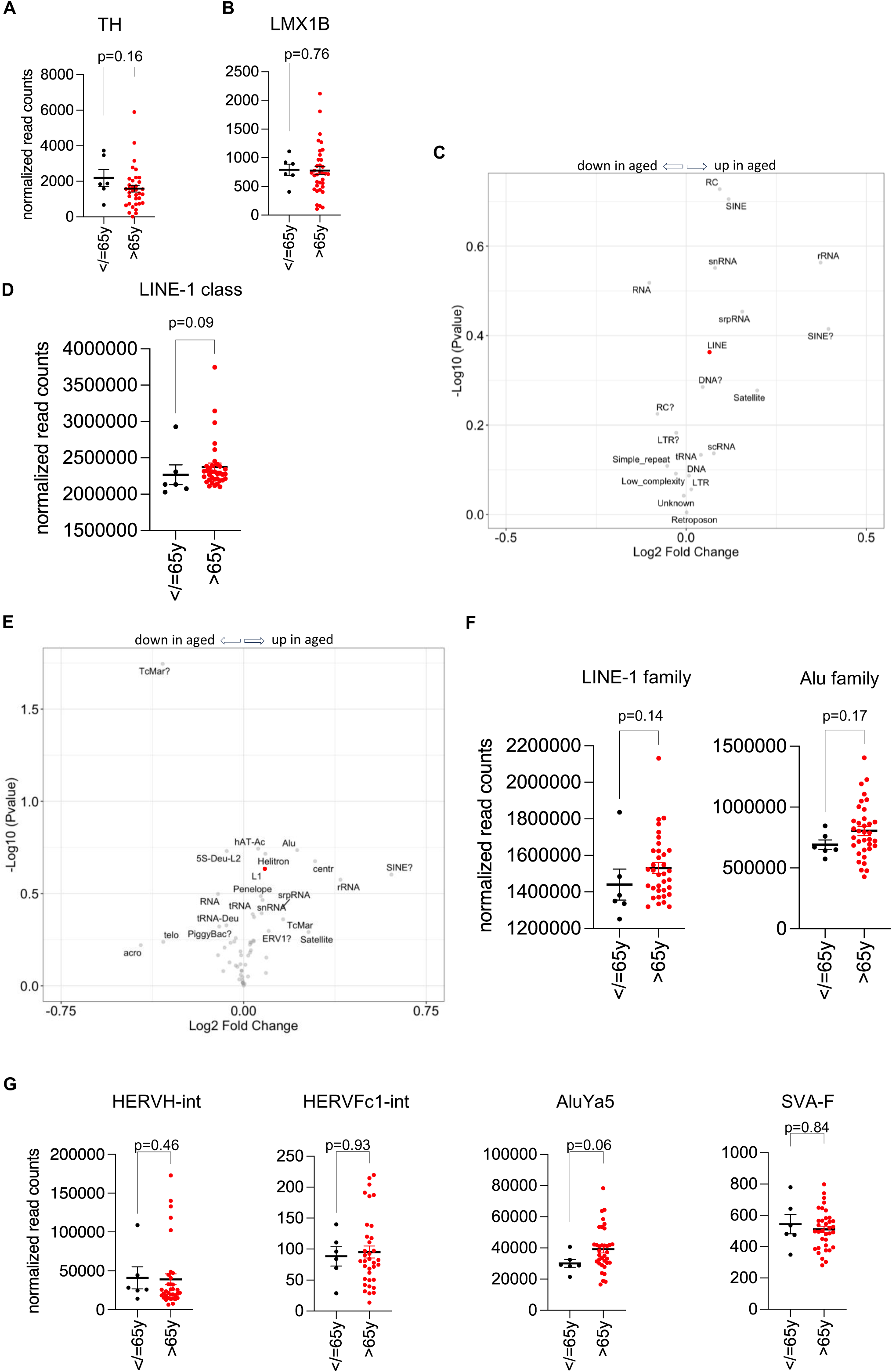
(A-B) Scatter plots comparing of the expression of dopaminergic markers tyrosine hydroxylase (TH, A) and LMX1B (B) between young (≤65y, n=6)) and aged (>65y, n=35) human dopaminergic neurons; Mann Whitney test. (C, E) Volcano plot of differential expression analysis of TE expression using DEseq2 comparing young (≤65y, n=6) and aged (>65y, n=35) human dopaminergic neurons at the “class” (C) and “family” (E) level of RepeatMasker. (D) Scatter plot comparing the expression of LINE at the “class” level between young (≤65y, n=6)) and aged (>65y, n=35) human dopaminergic neurons; Mann Whitney test. (F) Scatter plot comparing the expression of LINE and Alu at the “family” level between young (≤65y, n=6)) and aged (>65y, n=35) human dopaminergic neurons; Mann Whitney test. (G) Scatter plot comparing the expression of HERVH-int, HERV-Fc1 and two non-coding, non-autonomous but active TEs in the human genome, AluYa5 and SVA-F at the “name” level between young (≤65y, n=6)) and aged (>65y, n=35) human dopaminergic neurons; Mann Whitney test.

**Suppl Fig. 7.**
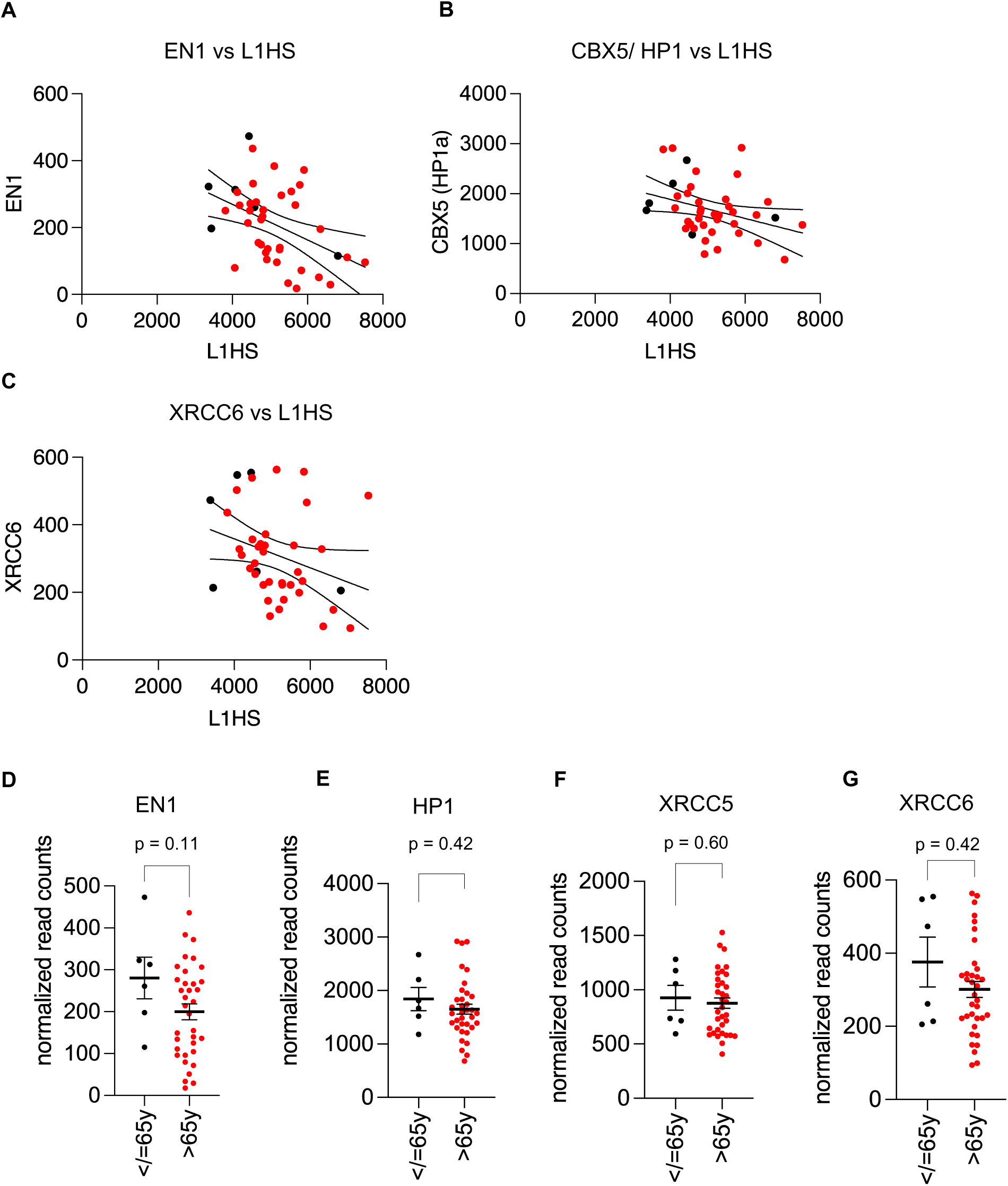
(A-C) Correlation analyses of L1HS expression with Engrailed 1 (EN1, A, Spearman r=-0.43, p=0.002), CBX5/HP1 (B, Spearman r=-0.35, p=0.01) and XRCC6 expression (C, Spearman r= -0.394, p=0.005). Normalized read counts are plotted. Black dots correspond to young individuals (≤65y), red dots correspond to aged individuals (>65y). (D-G) Scatter plots comparing the expression of EN1, CBX5/HP1, XRCC5 and XRCC6 between young (≤65y, n=6)) and aged (>65y, n=35) human dopaminergic neurons. Student’s t-test (EN1) or Mann Whitney test (HP1, XRCC5/6).

**Suppl Fig. 8.**
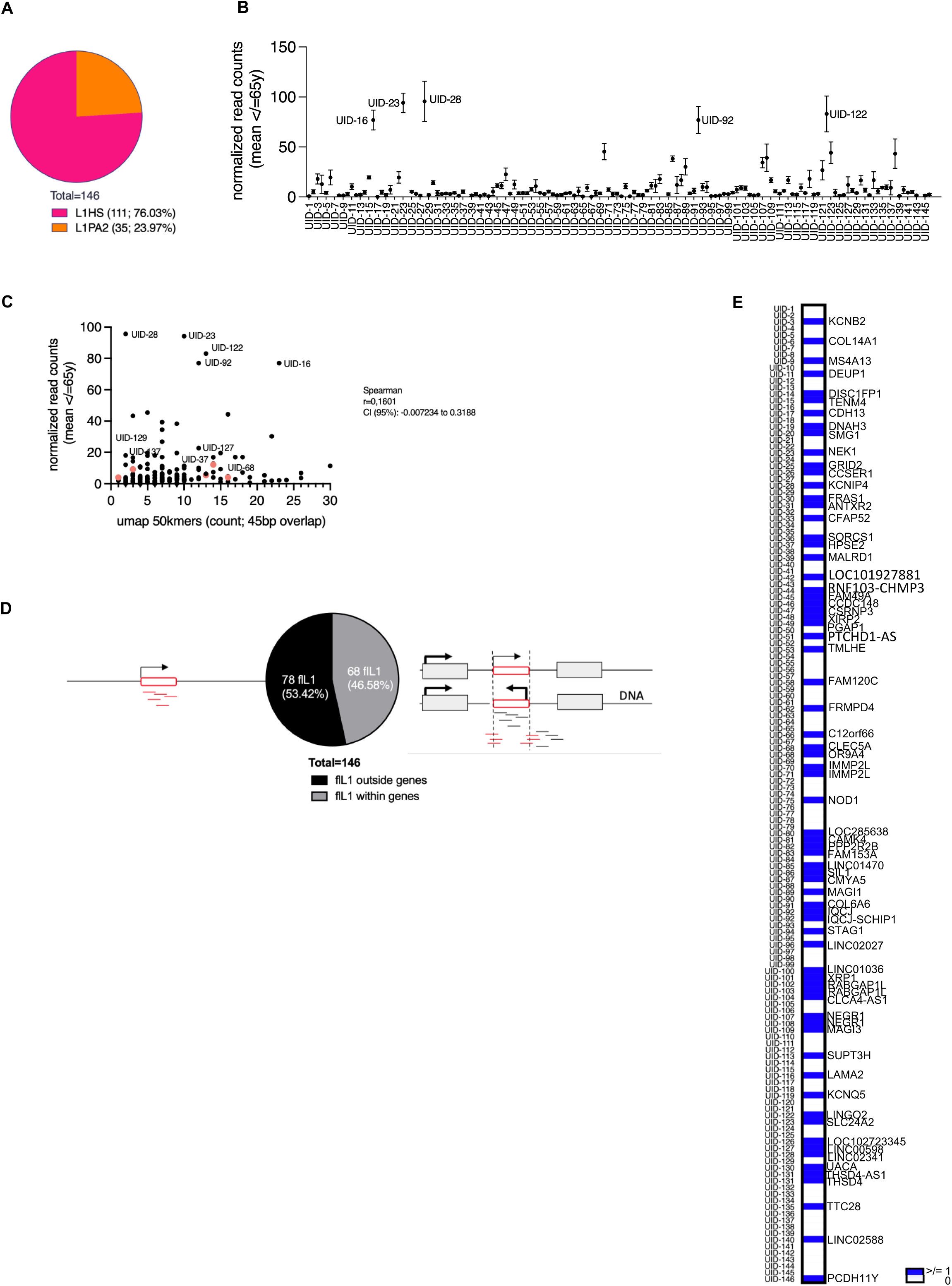
(B) *In silico* analysis of annotated full-length LINE-1 elements as in L1Basev2 (human reference genome hg38). Percentage of L1HS and L1PA2 elements among the 146 full-length elements (UID1-146). (C) Mean expression of all 146 full-length LINE-1 elements in dopaminergic neurons of all individuals ≤65y. (D) Correlation of mappability (UMAP hit counts over UID, see methods) and UID expression (normalized read counts). Spearman correlation. (E) *In silico* analysis of annotated full-length LINE-1 elements as in L1Basev2 (human reference genome hg38). Percentage of full-length LINE-1 elements located inside or outside a gene. (F) *In silico* analysis of annotated full-length LINE-1 elements as in L1Basev2 (human reference genome hg38). Presence (blue, with gene symbol) or absence (white) of a “hosting” gene among the 146 annotated full-length LINE-1 in the human reference genome.

## References

1. Lu, S. et al. A hidden human proteome encoded by ‘non-coding’ genes. Nucleic Acids Res 47, 8111–8125 (2019).

2. Lander, E. S. et al. Initial sequencing and analysis of the human genome. Nature 409, 860–921 (2001).

3. Brouha, B. et al. Hot L1s account for the bulk of retrotransposition in the human population. Proceedings of the National Academy of Sciences of the United States of America 100, 5280–5285 (2003).

4. Penzkofer, T. et al. L1Base 2: more retrotransposition-active LINE-1s, more mammalian genomes. Nucleic Acids Research 45, D68–D73 (2017).

5. Mao, J., Zhang, Q. & Cong, Y.-S. Human endogenous retroviruses in development and disease. Computational and Structural Biotechnology Journal 19, 5978–5986 (2021).

6. Esnault, C., Maestre, J. & Heidmann, T. Human LINE retrotransposons generate processed pseudogenes. Nat Genet 24, 363–367 (2000).

7. Wei, W. et al. Human L1 retrotransposition: cis preference versus trans complementation. Molecular and Cellular Biology 21, 1429–1439 (2001).

8. Mathias, S. L., Scott, A. F., Kazazian, H. H., Boeke, J. D. & Gabriel, A. Reverse transcriptase encoded by a human transposable element. Science 254, 1808–1810 (1991).

9. Feng, Q., Moran, J. V., Kazazian, H. H. & Boeke, J. D. Human L1 retrotransposon encodes a conserved endonuclease required for retrotransposition. Cell 87, 905–916 (1996).

10. Chen, L., Dahlstrom, J. E., Lee, S.-H. & Rangasamy, D. Naturally occurring endo-siRNA silences LINE-1 retrotransposons in human cells through DNA methylation. Epigenetics : official journal of the DNA Methylation Society 7, 758–771 (2012).

11. Goodier, J. L., Cheung, L. E. & Kazazian, H. H. MOV10 RNA helicase is a potent inhibitor of retrotransposition in cells. PLoS Genetics 8, e1002941 (2012).

12. Philippe, C. et al. Activation of individual L1 retrotransposon instances is restricted to cell-type dependent permissive loci. eLife 5, (2016).

13. Peze-Heidsieck, E. et al. Retrotransposons as a Source of DNA Damage in Neurodegeneration. Frontiers in aging neuroscience 13, 786897 (2021).

14. Driver, C. J. & McKechnie, S. W. Transposable elements as a factor in the aging of Drosophila melanogaster. Annals of the New York Academy of Sciences 673, 83–91 (1992).

15. Copley, K. E. & Shorter, J. Repetitive elements in aging and neurodegeneration. Trends in Genetics S0168952523000343 (2023) doi:10.1016/j.tig.2023.02.008.

16. Gorbunova, V. et al. The role of retrotransposable elements in ageing and age-associated diseases. Nature 1–11 (2021) doi:10.1038/s41586-021-03542-y.

17. Blaudin de Thé, F.-X., et al. Engrailed homeoprotein blocks degeneration in adult dopaminergic neurons through LINE-1 repression. The EMBO Journal 37, (2018).

18. Bodea, G. O. et al. LINE-1 retrotransposons contribute to mouse PV interneuron development. Nat Neurosci (2024) doi:10.1038/s41593-024-01650-2.

19. Sur, D. et al. Detection of the LINE-1 retrotransposonRNA-binding protein ORF1p in differentanatomical regions of the human brain. Mobile DNA 1–12 (2017) doi:10.1186/s13100-017-0101-4.

20. McKerrow, W. et al. LINE-1 retrotransposon expression in cancerous, epithelial and neuronal cells revealed by 5′ single-cell RNA-Seq. Nucleic Acids Research 51, 2033–2045 (2023).

21. Gasior, S. L., Wakeman, T. P., Xu, B. & Deininger, P. L. The human LINE-1 retrotransposon creates DNA double-strand breaks. Journal of molecular biology 357, 1383– 1393 (2006).

22. Belgnaoui, S. M., Gosden, R. G., Semmes, O. J. & Haoudi, A. Human LINE-1 retrotransposon induces DNA damage and apoptosis in cancer cells. Cancer cell international 6, 13 (2006).

23. De Cecco, M. et al. Genomes of replicatively senescent cells undergo global epigenetic changes leading to gene silencing and activation of transposable elements. Aging Cell 12, 247–256 (2013).

24. Van Meter, M. et al. SIRT6 represses LINE1 retrotransposons by ribosylating KAP1 but this repression fails with stress and age. Nature communications 5, 5011 (2014).

25. Sturm, Á., Ivics, Z. & Vellai, T. The mechanism of ageing: primary role of transposable elements in genome disintegration. Cell. Mol. Life Sci. 72, 1839–1847 (2015).

26. Wallace, N., Wagstaff, B. J., Deininger, P. L. & Roy-Engel, A. M. LINE-1 ORF1 protein enhances Alu SINE retrotransposition. Gene 419, 1–6 (2008).

27. Thomas, C. A. et al. Modeling of TREX1-Dependent Autoimmune Disease using Human Stem Cells Highlights L1 Accumulation as a Source of Neuroinflammation. Cell stem cell 21, 319–331.e8 (2017).

28. De Cecco, M. et al. L1 drives IFN in senescent cells and promotes age-associated inflammation. Nature 566, 73–78 (2019).

29. Luqman-Fatah, A. et al. The interferon stimulated gene-encoded protein HELZ2 inhibits human LINE-1 retrotransposition and LINE-1 RNA-mediated type I interferon induction. Nat Commun 14, 203 (2023).

30. Gazquez-Gutierrez, A., Witteveldt, J., R Heras, S. & Macias, S. Sensing of transposable elements by the antiviral innate immune system. RNA (New York, N.Y.) (2021) doi:10.1261/rna.078721.121.

31. Krug, L. et al. Retrotransposon activation contributes to neurodegeneration in a Drosophila TDP-43 model of ALS. PLoS Genetics 13, e1006635 (2017).

32. Casale, A. M. et al. Transposable element activation promotes neurodegeneration in a Drosophila model of Huntington’s disease. ISCIENCE 25, 103702 (2022).

33. Takahashi, T. et al. LINE-1 activation in the cerebellum drives ataxia. Neuron 1–19 (2022) doi:10.1016/j.neuron.2022.08.011.

34. Chiaruttini, N. et al. ABBA+BraiAn, an integrated suite for whole-brain mapping, reveals brain-wide differences in immediate-early genes induction upon learning. Cell Rep 44, 115876 (2025).

35. De Luca, C., Gupta, A. & Bortvin, A. Retrotransposon LINE-1 bodies in the cytoplasm of piRNA-deficient mouse spermatocytes: Ribonucleoproteins overcoming the integrated stress response. PLoS Genet 19, e1010797 (2023).

36. Spencley, A. L. et al. Co-transcriptional genome surveillance by HUSH is coupled to termination machinery. Mol Cell 83, 1623–1639.e8 (2023).

37. Garland, W. et al. Chromatin modifier HUSH co-operates with RNA decay factor NEXT to restrict transposable element expression. Mol Cell 82, 1691–1707.e8 (2022).

38. Shirane, K., Miura, F., Ito, T. & Lorincz, M. C. NSD1-deposited H3K36me2 directs de novo methylation in the mouse male germline and counteracts Polycomb-associated silencing. Nat Genet 52, 1088–1098 (2020).

39. Guerrero, A. et al. Cardiac glycosides are broad-spectrum senolytics. Nat Metab 1, 1074–1088 (2019).

40. Sato, S. et al. LINE-1 ORF1p as a candidate biomarker in high grade serous ovarian carcinoma. Sci Rep 13, 1537 (2023).

41. McKerrow, W. et al. LINE-1 expression in cancer correlates with p53 mutation, copy number alteration, and S phase checkpoint. Proc Natl Acad Sci U S A 119, e2115999119 (2022).

42. Walter, M., Teissandier, A., Pérez-Palacios, R. & Bourc’his, D. An epigenetic switch ensures transposon repression upon dynamic loss of DNA methylation in embryonic stem cells. eLife 5, e11418 (2016).

43. Larson, P. A. et al. Spliced integrated retrotransposed element (SpIRE) formation in the human genome. PLoS Biol 16, e2003067 (2018).

44. Müller, I. et al. MPP8 is essential for sustaining self-renewal of ground-state pluripotent stem cells. Nat Commun 12, 3034 (2021).

45. Znaidi, R. et al. Nuclear translocation of the LINE-1 encoded ORF1 protein alters nuclear envelope integrity in human neurons. Brain Res 1857, 149579 (2025).

46. Gusel’nikova, V. V. & Korzhevskiy, D. E. NeuN As a Neuronal Nuclear Antigen and Neuron Differentiation Marker. Acta Naturae 7, 42–47 (2015).

47. Cannon, J. R. & Greenamyre, J. T. NeuN is not a reliable marker of dopamine neurons in rat substantia nigra. Neuroscience Letters 464, 14–17 (2009).

48. Sukhorukova, E. G. Nuclear Protein NeuN in Neurons in the Human Substantia Nigra. Neurosci Behav Physi 44, 539–541 (2014).

49. Kumar, S. S. & Buckmaster, P. S. Neuron-specific nuclear antigen NeuN is not detectable in gerbil subtantia nigra pars reticulata. Brain Res 1142, 54–60 (2007).

50. Keller, D., Erö, C. & Markram, H. Cell Densities in the Mouse Brain: A Systematic Review. Front. Neuroanat. 12, 83 (2018).

51. Sun, W. et al. SOX9 Is an Astrocyte-Specific Nuclear Marker in the Adult Brain Outside the Neurogenic Regions. J Neurosci 37, 4493–4507 (2017).

52. Yushkova, E. & Moskalev, A. Transposable elements and their role in aging. Ageing Res Rev 86, 101881 (2023).

53. Gibb, W. R. & Lees, A. J. Anatomy, pigmentation, ventral and dorsal subpopulations of the substantia nigra, and differential cell death in Parkinson’s disease. J Neurol Neurosurg Psychiatry 54, 388–396 (1991).

54. Dong, X. et al. Enhancers active in dopamine neurons are a primary link between genetic variation and neuropsychiatric disease. Nature Neuroscience 1–19 (2018) doi:10.1038/s41593-018-0223-0.

55. Teissandier, A., Servant, N., Barillot, E. & Bourc’his, D. Tools and best practices for retrotransposon analysis using high-throughput sequencing data. Mobile DNA 10, 52 (2019).

56. Liu, N. et al. Selective silencing of euchromatic L1s revealed by genome-wide screens for L1 regulators. Nature 553, 228–232 (2018).

57. Rekaik, H., Blaudin de Thé, F.-X., Prochiantz, A., Fuchs, J. & Joshi, R. L. Dissecting the role of Engrailed in adult dopaminergic neurons–Insights into Parkinson disease pathogenesis. FEBS Letters 589, 3786–3794 (2015).

58. Maeda, R. & Tachibana, M. HP1 maintains protein stability of H3K9 methyltransferases and demethylases. EMBO reports 23, e53581 (2022).

59. Suzuki, J. et al. Genetic Evidence That the Non-Homologous End-Joining Repair Pathway Is Involved in LINE Retrotransposition. PLoS Genetics 5, e1000461 (2009).

60. Goerner-Potvin, P. & Bourque, G. Computational tools to unmask transposable elements. Nature Reviews Genetics 1–17 (2018) doi:10.1038/s41576-018-0050-x.

61. Schwarz, R., Koch, P., Wilbrandt, J. & Hoffmann, S. Locus-specific expression analysis of transposable elements. Briefings in Bioinformatics 23, bbab417 (2022).

62. Faulkner, G. J. Elevated L1 expression in ataxia telangiectasia likely explained by an RNA-seq batch effect. Neuron 111, 610–611 (2023).

63. Lanciano, S. & Cristofari, G. Measuring and interpreting transposable element expression. Nature Reviews Genetics 509, 7 (2020).

64. Martin, S. L., Li, J. & Weisz, J. A. Deletion analysis defines distinct functional domains for protein-protein and nucleic acid interactions in the ORF1 protein of mouse LINE-1. J Mol Biol 304, 11–20 (2000).

65. Briggs, E. M. et al. RIP-seq reveals LINE-1 ORF1p association with p-body enriched mRNAs. 1–13 (2021) doi:10.1186/s13100-021-00233-3.

66. Kulpa, D. A. & Moran, J. V. Ribonucleoprotein particle formation is necessary but not sufficient for LINE-1 retrotransposition. Human Molecular Genetics 14, 3237–3248 (2005).

67. Taylor, M. S. et al. Dissection of affinity captured LINE-1 macromolecular complexes. eLife 7, 210–41 (2018).

68. Kelly, M. P. et al. Select 3’,5’-cyclic nucleotide phosphodiesterases exhibit altered expression in the aged rodent brain. Cell Signal 26, 383–397 (2014).

69. Lakics, V., Karran, E. H. & Boess, F. G. Quantitative comparison of phosphodiesterase mRNA distribution in human brain and peripheral tissues. Neuropharmacology 59, 367–374 (2010).

70. Mandel, S. & Gozes, I. Activity-dependent neuroprotective protein constitutes a novel element in the SWI/SNF chromatin remodeling complex. J Biol Chem 282, 34448–34456 (2007).

71. Singh, A., Modak, S. B., Chaturvedi, M. M. & Purohit, J. S. SWI/SNF Chromatin Remodelers: Structural, Functional and Mechanistic Implications. Cell Biochem Biophys 81, 167–187 (2023).

72. Goodier, J. L., Cheung, L. E. & Kazazian, H. H. Mapping the LINE1 ORF1 protein interactome reveals associated inhibitors of human retrotransposition. Nucleic Acids Research 41, 7401–7419 (2013).

73. Taylor, M. S. et al. Affinity proteomics reveals human host factors implicated in discrete stages of LINE-1 retrotransposition. Cell 155, 1034–1048 (2013).

74. Moldovan, J. B. & Moran, J. V. The Zinc-Finger Antiviral Protein ZAP Inhibits LINE and Alu Retrotransposition. PLoS Genetics 11, e1005121 (2015).

75. Vuong, L. M., Pan, S. & Donovan, P. J. Proteome Profile of Endogenous Retrotransposon-Associated Complexes in Human Embryonic Stem Cells. Proteomics 19, e1900169 (2019).

76. Ardeljan, D. et al. LINE-1 ORF2p expression is nearly imperceptible in human cancers. Mobile DNA 11, 1 (2020).

77. Liu, Q. et al. MOV10 recruits DCP2 to decap human LINE-1 RNA by forming large cytoplasmic granules with phase separation properties. EMBO reports 24, e56512 (2023).

78. Rodić, N. et al. Long Interspersed Element-1 Protein Expression Is a Hallmark of Many Human Cancers. Am J Pathol 184, 1280–1286 (2014).

79. Osburn, S. C. et al. Long-term voluntary wheel running effects on markers of long interspersed nuclear element-1 in skeletal muscle, liver, and brain tissue of female rats. American Journal of Physiology-Cell Physiology 323, C907–C919 (2022).

80. Rybacki, K., Xia, M., Ahsan, M. U., Xing, J. & Wang, K. Assessing the Expression of Long INterspersed Elements (LINEs) via Long-Read Sequencing in Diverse Human Tissues and Cell Lines. Genes 14, 1893 (2023).

81. Doucet-O’Hare, T. T. et al. LINE-1 expression and retrotransposition in Barrett’s esophagus and esophageal carcinoma. Proceedings of the National Academy of Sciences 112, E4894–E4900 (2015).

82. Goodier, J. L., Ostertag, E. M., Du, K. & Kazazian, H. H. A Novel Active L1 Retrotransposon Subfamily in the Mouse. Genome Res. 11, 1677–1685 (2001).

83. Bodak, M., Yu, J. & Ciaudo, C. Regulation of LINE-1 in mammals. Biomolecular Concepts 5, 409–428 (2014).

84. Naas, T. P. et al. An actively retrotransposing, novel subfamily of mouse L1 elements. EMBO J 17, 590–597 (1998).

85. Kuwabara, T. et al. Wnt-mediated activation of NeuroD1 and retro-elements during adult neurogenesis. Nat Neurosci 12, 1097–1105 (2009).

86. Zhang, Y. et al. Single-cell epigenome analysis reveals age-associated decay of heterochromatin domains in excitatory neurons in the mouse brain. Cell Res 32, 1008–1021 (2022).

87. Garza, R. et al. LINE-1 retrotransposons drive human neuronal transcriptome complexity and functional diversification. Sci Adv 9, eadh9543 (2023).

88. Watanabe, R. et al. Identification of epigenetically active L1 promoters in the human brain and their relationship with psychiatric disorders. Neuroscience Research S0168010223000913 (2023) doi:10.1016/j.neures.2023.05.001.

89. Takahashi, F. et al. Immune-mediated neurodegenerative trait provoked by multimodal derepression of long-interspersed nuclear element-1. iScience 25, 104278 (2022).

90. Simon, M. et al. LINE1 Derepression in Aged Wild-Type and SIRT6-Deficient Mice Drives Inflammation. Cell metabolism 29, 871–885.e5 (2019).

91. Lopez-Otín, C., Blasco, M. A., Partridge, L., Serrano, M. & Kroemer, G. Hallmarks of aging: An expanding universe. Cell S0092867422013770 (2023) doi:10.1016/j.cell.2022.11.001.

92. Mumford, P. W. et al. Skeletal muscle LINE-1 retrotransposon activity is upregulated in older versus younger rats. Am J Physiol Regul Integr Comp Physiol 317, R397–R406 (2019).

93. Dai, L., LaCava, J., Taylor, M. S. & Boeke, J. D. Expression and detection of LINE-1 ORF-encoded proteins. Mobile genetic elements 4, e29319 (2014).

94. Whongsiri, P. et al. Many Different LINE-1 Retroelements Are Activated in Bladder Cancer. International Journal of Molecular Sciences 21, 9433–14 (2020).

95. Lindtner, S., Felber, B. K. & Kjems, J. An element in the 3’ untranslated region of human LINE-1 retrotransposon mRNA binds NXF1(TAP) and can function as a nuclear export element. RNA 8, 345–356 (2002).

96. Naufer, M. N., Furano, A. V. & Williams, M. C. Protein-nucleic acid interactions of LINE-1 ORF1p. Seminars in Cell & Developmental Biology 86, 140–149 (2019).

97. Garneau, N. L., Wilusz, J. & Wilusz, C. J. The highways and byways of mRNA decay. Nat Rev Mol Cell Biol 8, 113–126 (2007).

98. Lykke-Andersen, J. Identification of a human decapping complex associated with hUpf proteins in nonsense-mediated decay. Mol Cell Biol 22, 8114–8121 (2002).

99. Threlfell, S., Sammut, S., Menniti, F. S., Schmidt, C. J. & West, A. R. Inhibition of Phosphodiesterase 10A Increases the Responsiveness of Striatal Projection Neurons to Cortical Stimulation. J Pharmacol Exp Ther 328, 785–795 (2009).

100. Li, N. et al. Suppression of β-catenin/TCF transcriptional activity and colon tumor cell growth by dual inhibition of PDE5 and 10. Oncotarget 6, 27403–27415 (2015).

101. Pereira, G. C. et al. Properties of LINE-1 proteins and repeat element expression in the context of amyotrophic lateral sclerosis. Mobile DNA 9, 35 (2018).

102. Mita, P. et al. LINE-1 protein localization and functional dynamics during the cell cycle. eLife 7, (2018).

103. Newton, J. C. et al. Phase separation of the LINE-1 ORF1 protein is mediated by the N-terminus and coiled-coil domain. Biophysical journal 120, 2181–2191 (2021).

104. Linkert, M. et al. Metadata matters: access to image data in the real world. J Cell Biol 189, 777–782 (2010).

105. Ollion, J., Cochennec, J., Loll, F., Escudé, C. & Boudier, T. TANGO: a generic tool for high-throughput 3D image analysis for studying nuclear organization. *Bioinformatics (Oxford*, England) 29, 1840–1841 (2013).

106. Stringer, C., Wang, T., Michaelos, M. & Pachitariu, M. Cellpose: a generalist algorithm for cellular segmentation. Nat Methods 18, 100–106 (2021).

107. Wang, Q. et al. The Allen Mouse Brain Common Coordinate Framework: A 3D Reference Atlas. Cell 181, 936–953.e20 (2020).

108. Chiaruttini, N. et al. An Open-Source Whole Slide Image Registration Workflow at Cellular Precision Using Fiji, QuPath and Elastix. Front. Comput. Sci. 3, 780026 (2022).

109. Bankhead, P. et al. QuPath: Open source software for digital pathology image analysis. Sci Rep 7, 16878 (2017).

110. Schmidt, U., Weigert, M., Broaddus, C. & Myers, G. Cell Detection with Star-Convex Polygons. in Medical Image Computing and Computer Assisted Intervention – MICCAI 2018 (eds. Frangi, A. F., Schnabel, J. A., Davatzikos, C., Alberola-López, C. & Fichtinger, G.) 265–273 (Springer International Publishing, Cham, 2018). doi:10.1007/978-3-030-00934-2_30.

111. McKinney, W. Data Structures for Statistical Computing in Python. in 56–61 (Austin, Texas, 2010). doi:10.25080/Majora-92bf1922-00a.

112. Liao, Y., Smyth, G. K. & Shi, W. The R package Rsubread is easier, faster, cheaper and better for alignment and quantification of RNA sequencing reads. Nucleic Acids Res 47, e47 (2019).

113. Poullet, P., Carpentier, S. & Barillot, E. myProMS, a web server for management and validation of mass spectrometry-based proteomic data. Proteomics 7, 2553–2556 (2007).

114. Valot, B., Langella, O., Nano, E. & Zivy, M. MassChroQ: a versatile tool for mass spectrometry quantification. Proteomics 11, 3572–3577 (2011).

115. Perez-Riverol, Y. et al. The PRIDE database resources in 2022: a hub for mass spectrometry-based proteomics evidences. Nucleic Acids Res 50, D543–D552 (2022).

116. Thomas, P. D. et al. PANTHER: Making genome-scale phylogenetics accessible to all. Protein Sci 31, 8–22 (2022).

117. Shannon, P. et al. Cytoscape: a software environment for integrated models of biomolecular interaction networks. Genome Res 13, 2498–2504 (2003).

